# Genome-scale and pathway engineering for the sustainable aviation fuel precursor isoprenol production in *Pseudomonas putida*

**DOI:** 10.1101/2023.04.29.538800

**Authors:** Deepanwita Banerjee, Ian S. Yunus, Xi Wang, Jinho Kim, Aparajitha Srinivasan, Russel Menchavez, Yan Chen, Jennifer W. Gin, Christopher J. Petzold, Hector Garcia Martin, Paul D. Adams, Aindrila Mukhopadhyay, Joonhoon Kim, Taek Soon Lee

## Abstract

Sustainable aviation fuel (SAF) will significantly impact global warming in the aviation sector, and important SAF targets are emerging. Isoprenol is a precursor for a promising SAF compound DMCO (1,4-dimethylcyclooctane), and has been produced in several engineered microorganisms. Recently, *Pseudomonas putida* has gained interest as a future host for isoprenol bioproduction as it can utilize carbon sources from inexpensive plant biomass. Here, we engineer metabolically versatile host *P. putida* for isoprenol production. We employ two computational modeling approaches (Bilevel optimization and Constrained Minimal Cut Sets) to predict gene knockout targets and optimize the “IPP-bypass” pathway in *P. putida* to maximize isoprenol production. Altogether, the highest isoprenol production titer from *P. putida* was achieved at 3.5 g/L under fed-batch conditions. This combination of computational modeling and strain engineering on *P. putida* for an advanced biofuels production has vital significance in enabling a bioproduction process that can use renewable carbon streams.

## Introduction

Biological production of aviation fuels and their precursors from sustainable carbon sources stands to have a realistic impact on reducing CO_2_ emissions, an increasingly critical aspect of addressing climate change^1, 2^. For this reason, several sustainable aviation fuel (SAF) targets and their precursors are being proposed, which include not only traditional ethanol-based fuels^3^, but also novel high-energy multicyclic compounds possible via bioproduction, such as fuelimycin A^4^ and epi-isozizaene^5, 6^. One such important precursor is isoprenol (a.k.a 3-methylbut-3-en-1-ol). A commodity platform chemical and a vetted biogasoline^7^, isoprenol is also the precursor for the jet fuel 1,4-dimethyl cyclooctane (DMCO). Catalytic conversion of isoprenol to DMCO has been shown at high efficiency^8^ and establishing a carbon-efficient conversion of renewable carbon sources to isoprenol would enable a highly sustainable process^8^ for this SAF.

While isoprenol production has been shown in model microbial hosts (*Escherichia coli*^9^, *Corynebacterium glutamicum*^10^, and *Saccharomyces cerevisiae*^11^), catabolically versatile microbes that consume a wider range of carbon compounds are essential to providing a cost-effective process^1, 12^. In the case of plant biomass conversion, there is an urgent need to demonstrate production of isoprenol in microbial systems that can catabolize both sugars and aromatics derived from lignocellulosic biomass. *Pseudomonas putida* KT2440 is an ideal conversion host with a versatile conversion profile^13, 14^ and efficient genetic tools. While prior works in model organisms^15, 16^ have achieved robust isoprenol titers, a microbial host such as *P. putida* KT2440 is a more likely candidate for the final deployment for isoprenol production as a SAF precursor. A less-commonly used laboratory host such as *P. putida*, however, is a far more challenging system to develop as a conversion platform. For instance, the most efficient route to isoprenol is through the heterologous mevalonate (MVA) pathway using an IPP-bypass that utilizes hydroxymethylglutaryl CoA (HMG-CoA) as the precursor^15^. However, efforts in *P. putida*^17^ have shown that the mere MVA pathway overexpression did not provide any improvements over the native 2-C-methyl-D-erythritol 4-phosphate (MEP) pathway overexpression and both resulted in very low titers. A similar observation was also reported in cyanobacteria^18^.

In a recent study, we were able to establish the MVA pathway in *P. putida* KT2440^19^ and define the necessary cultivation conditions to produce isoprenol in this host via the heterologous pathway. While this provides an excellent foundation for isoprenol production in *P. putida* KT2440, this microbes’ unusual metabolic profile presents several challenges that need to be overcome. One issue is the catabolism of isoprenol itself, and its intermediates, by *P. putida* KT2440. Extensive functional genomics data have recently been accumulated for *P. putida* KT2440 and have revealed genes associated with degradation or catabolism of non-canonical carbon sources (e.g., levulinic acid^20^, lysine^21^, and isoprenol^22^), and also provided the hypotheses for host engineering targets to optimize the desired catabolism and minimize the undesired ones.

Computationally driven metabolic engineering methods have gained interest during the last decade^23^. Such methods can predict strategies that may involve large numbers of genetic interventions (e.g. deletion, overexpression, or repression) to reach the predicted yields. Implementation of such strategies is sometimes challenging even with recent advances in synthetic biology and metabolic engineering techniques. To address this challenge and cover a larger solution space we used multiple computational strain design methods based on elementary mode analysis or bilevel optimization. The latest highly curated genome-scale metabolic model (GSMM) for *P. putida*^24^ enabled the use of these approaches and also highlighted the differences in metabolism from model microbes such as *E. coli*.

In this work, we employ two GSMM-guided approaches in combination with targeted edits and pathway improvements to enhance the production of the DMCO precursor, isoprenol, in *P. putida* KT2440 (**Figure 1**). We first add the knowledge from functional genomics data sets (e.g., genes involved in isoprenol degradation) and the heterologous MVA pathway to update the GSMM (**Figure 2**). We then use both Elementary mode analysis (EMA)-based methods^25, 26^ and bilevel optimization (Opt)-based methods^27, 28^ to prioritize a subset of host genome targets that, when deleted, are predicted to enhance flux to isoprenol via the MVA route. We also use known edits such as deletion of the *pha* operon and other literature-based targets to further enhance isoprenol titers. Finally, we use proteomics to optimize the pathway configuration. Overall, our GSMM-guided approach allowed us to select and prioritize the intervention targets, and lead to over 3.5 g/L isoprenol from glucose in a minimal defined medium under fed-batch conditions. This titer is the highest reported for *P. putida* KT2440 and has vital significance in enabling a bioproduction process that can use renewable carbon streams as the starting material.

**Figure 1.**
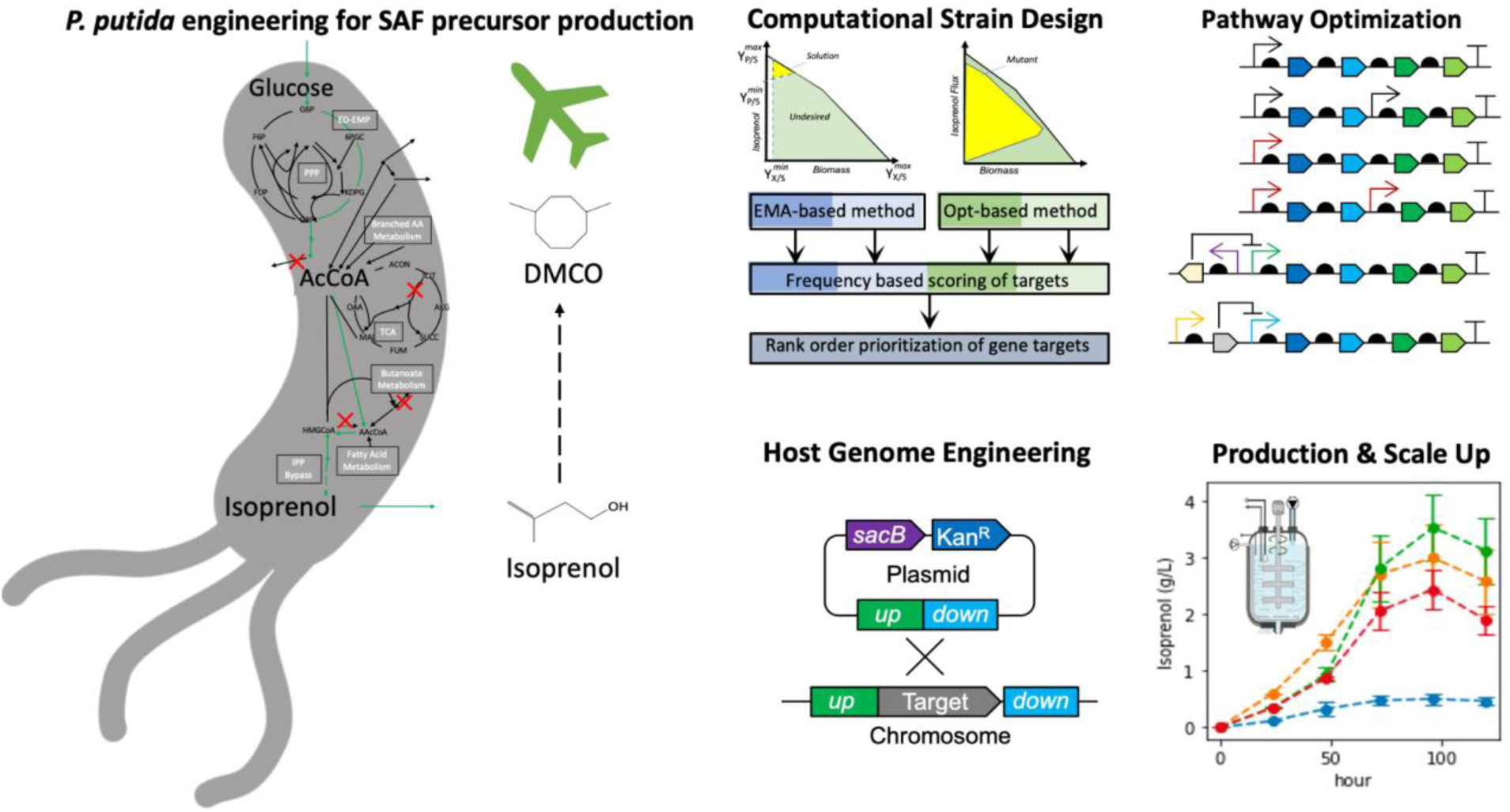
Genome-scale metabolic and pathway engineering for production of the sustainable aviation fuel (SAF) precursor, isoprenol, in *Pseudomonas putida*.

**Figure 2.**
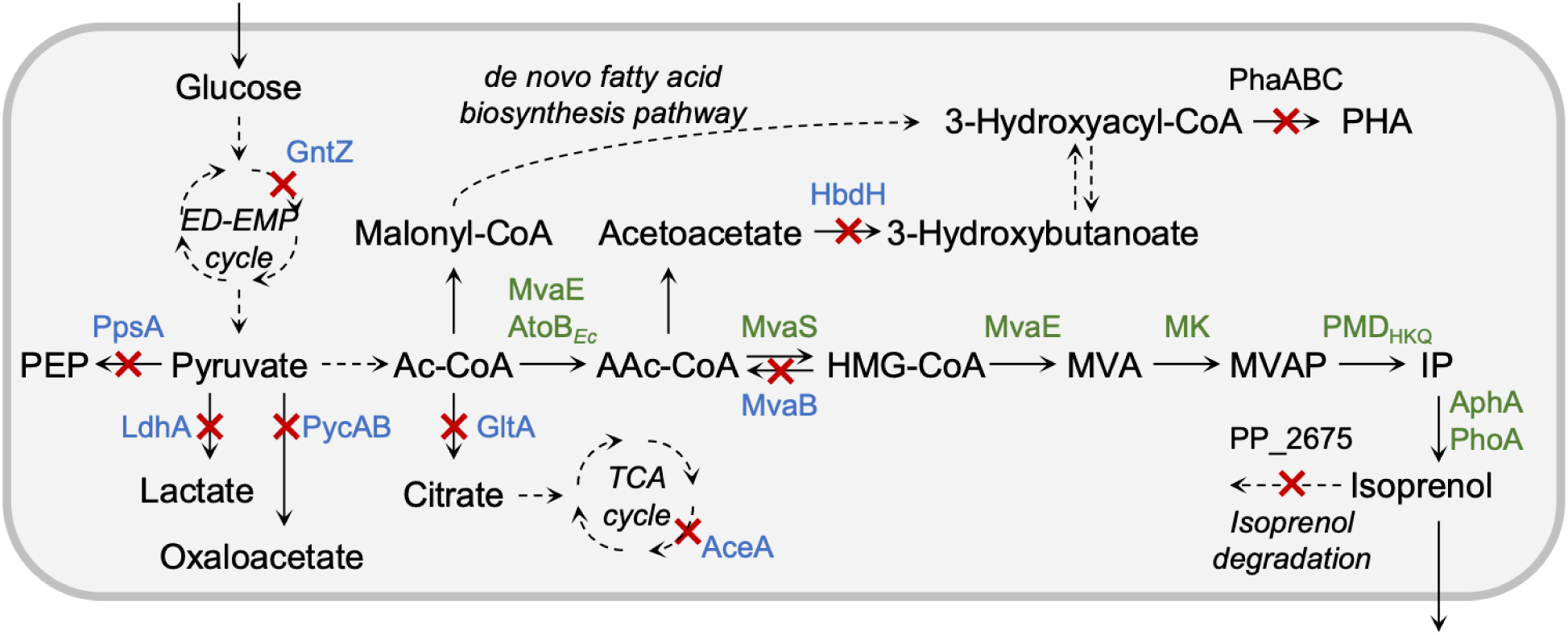
Metabolic pathways for isoprenol production using IPP bypass in *P. putida* KT2440. The gene knockout targets identified from genome-scale metabolic modeling are shown in blue. The heterologous genes for the IPP-bypass mevalonate pathway are shown in green.

## Results and Discussion

### Computational Strain Design for Isoprenol Production

For model-guided improvement of isoprenol production, we employed EMA-based approaches including Elementary Flux Modes (EFMs)^25^ and Constrained Minimal Cut Sets (cMCS)^26^ as well as Opt-based approaches including OptKnock^27, 29^ and OptForce^28^ using the latest GSMM for *P. putida* iJN1462^24^ augmented with the heterologous MVA pathway (<u>Supplementary File 1</u>). Preliminary computational strain design results from EMA-based and Opt-based methods showed that growth-coupled production of isoprenol requires the deletion of 9 or more metabolic reactions in *P. putida*. Construction of such mutants would require a large number of gene knockouts and significant experimental efforts, and intermediate mutants often need to be tested for growth and production to monitor the progress. However, it is not clear from computational predictions which genes are more important for increasing isoprenol production and therefore should be knocked out with high priority since the growth-coupled production does not happen *in silico* with the deletion of a subset of identified reactions. To this end, we generated a large number of computational designs using EMA-based and Opt-based methods, and calculated the frequency of engineering targets (i.e. fluxes to be shut down) appearing in the designs by each method. The frequency was used to calculate the rank order of targets for each design method, and the rank order from different methods was combined to calculate the final score. Our assumption was that certain targets can be more important for improving isoprenol production (e.g., due to higher fluxes or key branch points (or shunts)) than others and thus they will appear more frequently in a diverse set of computational designs. By generating a large number of designs using multiple computational methods and combining them using a rank-based ensemble approach, we aimed to identify such crucial targets and prioritize them for the experimental construction of knockout strains. Although the computational model requires the deletion of all targets from a design to see improved isoprenol production, we hypothesized that the deletion of a subset consisting of these crucial targets will still lead to improved isoprenol production.

For the EMA-based methods, we first calculated elementary flux modes (EFMs) using a small central metabolic network of *P. putida* with a lumped reaction that accounted for the heterologous isoprenol production pathway (<u>Supplementary File 2</u>). Each EFM is a minimal set of reactions carrying flux under the defined glucose minimal medium condition for growth as well as isoprenol production. A total of 360,475 EFMs were computed of which only 276 EFMs were selected that carried a flux through the biomass, ATP maintenance, glucose uptake, and isoprenol production reactions. These 276 EFMs showed an isoprenol yield ranging from 0.096 to 0.302 mol/mol of glucose and a biomass yield between 0.012 to 0.078. A frequency-based scoring was used to prioritize engineering targets, from 276 different computed EFMs (<u>Supplementary File 3</u>). Among the non-essential reactions that were predicted to carry no flux in majority of the EFMs included reactions LDH_D (100% of EFMs), PPS (100% of EFMs), PC (100% of EFMs), ICL/MALS (50% of the EFMs) and GND (26% of EFMs). Further we used cMCS to compute growth-coupled strategies for isoprenol production using glucose as the sole carbon source. From a total of 60 cMCS runs, we enumerated 4,950 feasible cMCS cut set designs. We used a frequency-based scoring to prioritize engineering targets from the different feasible cMCS designs that were computed for isoprenol and its precursors HMG-CoA, DMAPP or IPP (<u>Supplementary File 3</u>).

For the Opt-based methods, OptKnock was first used to find gene knockouts to couple isoprenol production to growth using the genome-scale metabolic model. Multiple solutions were identified by solving the OptKnock problem repeatedly and adding integer cuts. A total of 157 OptKnock solutions were initially collected and pre-processed to 120 solutions by removing solutions containing unnecessary or equivalent gene knockouts. In addition, we constrained the model by blocking the secretion of byproducts except for experimentally observed ones (e.g., gluconate, 2-ketogluconate, and acetate) and ran OptKnock to find another set of designs. Using the constrained model, a total of 377 OptKnock solutions were obtained and pre-processed to 263 solutions. OptForce was next used to identify strategies to improve isoprenol production. A total of 50 OptForce solutions were obtained, but we found that they consisted mostly of routes that increase or decrease flux and included only 9 potential knockout targets with low frequencies. Therefore, we decided to use the OptKnock solutions from two simulations to calculate the frequency for scoring gene targets (<u>Supplementary File 4</u>).

Finally, we combined scores from EMA-based (<u>Supplementary File 3</u>) and Opt-based (<u>Supplementary File 4</u>) predictions to arrive at the top 8 priority gene targets for experimental implementation (**Table 1 and Figure 2**). The first two priority gene targets (*mvaB* and *hbdH*) were involved in the degradation of endogenous metabolites (hydroxymethylglutaryl-CoA and acetoacetyl-CoA) that also participate in the heterologous MVA pathway. The other priority gene targets were involved in central carbon metabolism including the pentose phosphate pathway (*gntZ*), pyruvate metabolism (*ldhA*, *ppsA*, and *pycAB*), and TCA cycle (*gltA* and *aceA*).

**Table 1:**
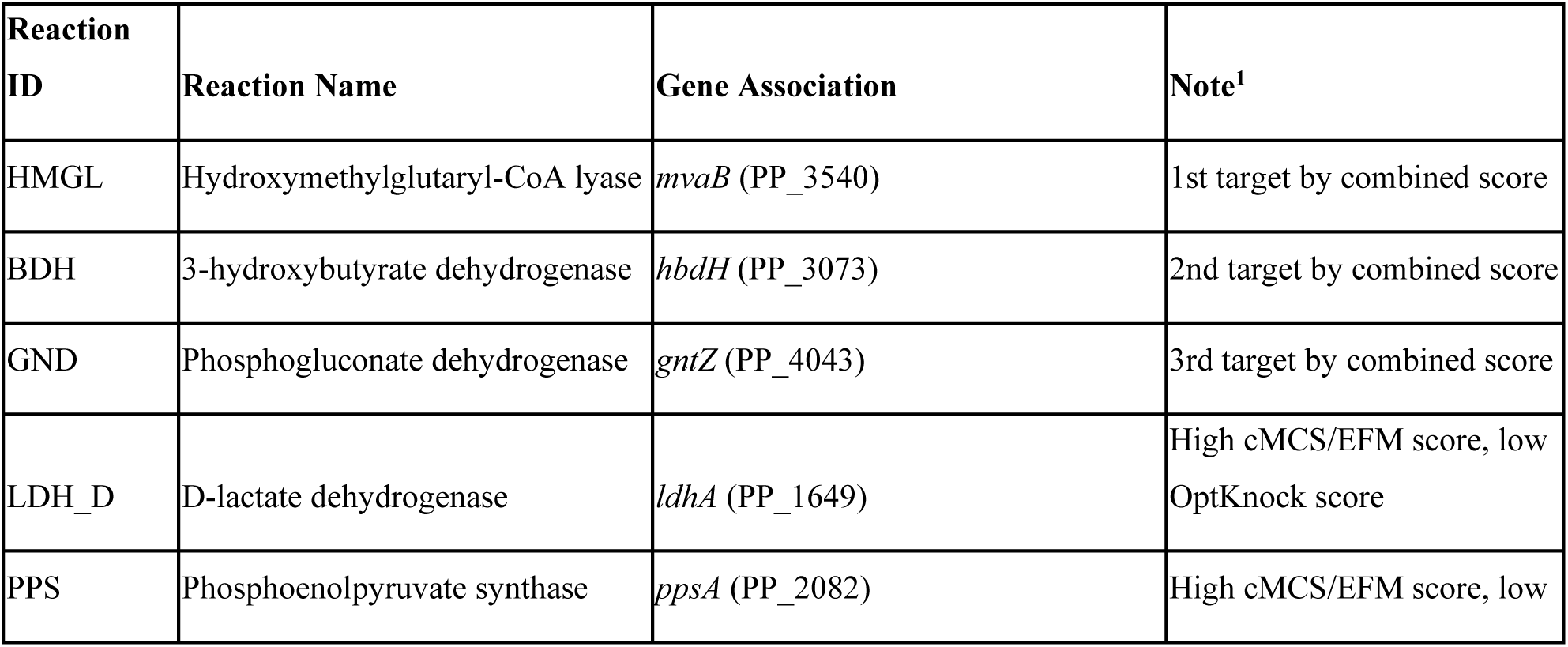
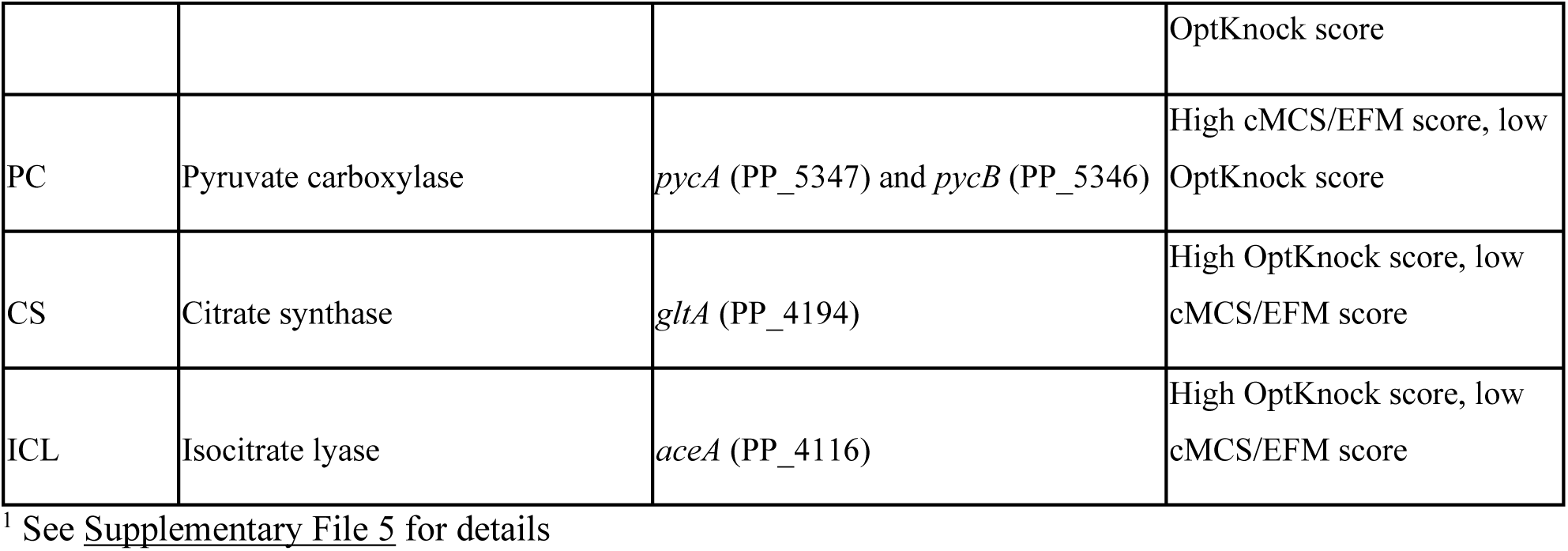
Priority Targets based on EMA-based and Opt-based approaches for experimental validation

| Reaction ID | Reaction Name | Gene Association | Note<sup>1</sup> |
| --- | --- | --- | --- |
| HMGL | Hydroxymethylglutaryl-CoA lyase | *mvaB* (PP_3540) | 1st target by combined score |
| BDH | 3-hydroxybutyrate dehydrogenase | *hbdH* (PP_3073) | 2nd target by combined score |
| GND | Phosphogluconate dehydrogenase | *gntZ* (PP_4043) | 3rd target by combined score |
| LDH_D | D-lactate dehydrogenase | *ldhA* (PP_1649) | High cMCS/EFM score, low OptKnock score |
| PPS | Phosphoenolpyruvate synthase | *ppsA* (PP_2082) | High cMCS/EFM score, low |
|  |  |  | OptKnock score |
| PC | Pyruvate carboxylase | *pycA* (PP_5347) and *pycB* (PP_5346) | High cMCS/EFM score, low OptKnock score |
| CS | Citrate synthase | *gltA* (PP_4194) | High OptKnock score, low cMCS/EFM score |
| ICL | Isocitrate lyase | *aceA* (PP_4116) | High OptKnock score, low cMCS/EFM score |

<sup>1</sup> See Supplementary File 5 for details

### Experimental Implementation of Metabolic Rewiring for Isoprenol Production

To experimentally verify the predicted gene knockout targets, the *P. putida* Δ*phaABC* strain (XW01, see **Supplementary Table 1** for the list of strains) was used as the background strain to perform the gene knockouts. This strain has shown the highest isoprenol level in *P. putida* (104 mg/L, XW11 strain) when using an IPP-bypass MVA pathway via a plasmid (pXW1, see **Supplementary Table 2** for the list of plasmids) in our previous work^19^. According to the core gene targets selected by multiple algorithms (**Table 1**), we constructed single and multiple knockout *P. putida* mutant strains to validate the model predictions (**Figure 3a**). We started the knockout set with the PP_3540/*mvaB* gene as it encodes for hydroxymethylglutaryl-CoA lyase which catalyzes a reaction transforming HMG-CoA into acetoacetate and acetyl-CoA, hence competing for HMG-CoA, a key precursor in mevalonate synthesis. The knockout of *mvaB* improved isoprenol production up to 164 mg/L, which increased 1.6-fold compared with the XW11 strain (the starter strain, **Figure 3b**).

**Figure 3.**
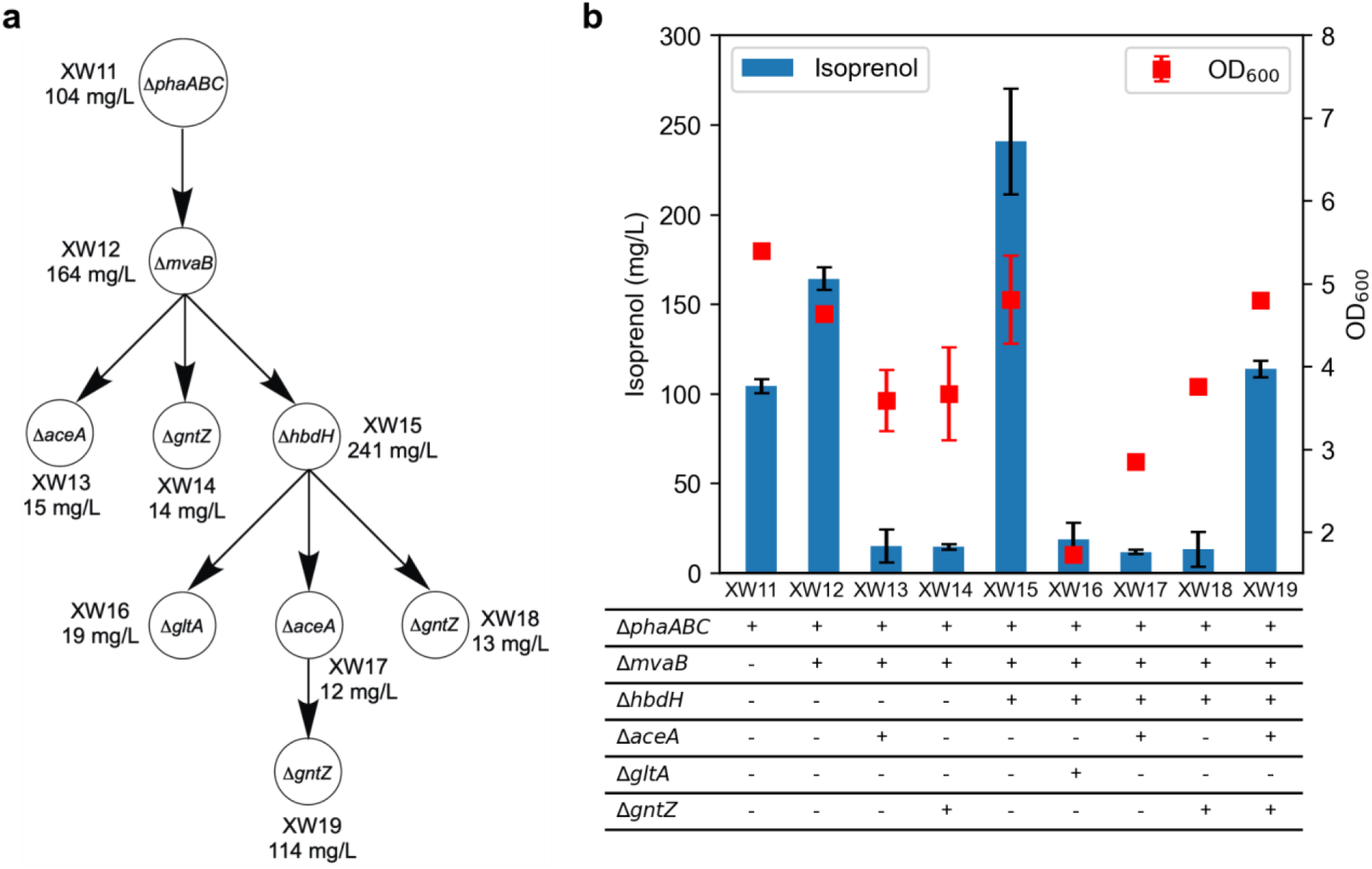
Engineering gene knockout mutants for experimental validation of genome-scale metabolic modeling predictions. **a** The flowchart of gene knockout strain development; **b** Isoprenol production and cell growth of constructed gene knockout strains after 48 hours using EZ rich medium. Data was obtained from three biological replicates and error bars represent standard deviation.

Using this double-knockout strain (Δ*phaABC* Δ*mvaB*) as a base, we performed a second round of gene knockouts with PP_4116/*aceA* (isocitrate lyase), PP_4043/*gntZ* (6-phosphogluconate dehydrogenase), and PP_3073/*hbdH* (3-hydroxybutyrate dehydrogenase) as targets (**Figure 3**). The knockout of *aceA* (XW13) or *gntZ* (XW14) significantly decreased isoprenol production to 14-15 mg/L, while the *hbdH* knockout (XW15) improved isoprenol production up to 241 mg/L, a 2.3-fold increase from the XW11 strain and 1.5-fold increase from the XW12 strain, respectively. As 3-hydroxybutyrate dehydrogenase (*hbdH*) catalyzes the conversion between 3-hydroxybutyrate and acetoacetate, the success of *hbdH* knockout in increasing isoprenol production might be attributed to the removal of a competing pathway of acetoacetyl-CoA, the first metabolite in the MVA pathway. Isocitrate lyase (*aceA*) converts isocitrate to glyoxylate, and 6-phosphogluconate dehydrogenase (*gntZ*) is a key enzyme at the shunt of the pentose phosphate pathway and the ED pathway. The failure of these two knockouts in isoprenol improvement might be attributed to the fact that they are both involved in central carbon metabolism, which is more robust than a specific biosynthesis pathway, such as the MVA pathway and its related metabolites. While Δ*aceA* or Δ*gntZ* did not improve isoprenol production, it is important to note that the algorithms we used in genome-scale metabolic modeling predicted the optimal production yield for combinations of gene knockouts, rather than a single gene deletion. Thus, engineering multiple gene knockouts in one strain may be still required to push the experimental results closer to the modeling prediction.

Therefore, for the third round of knockouts, we picked the highest producer with the triple knockouts (Δ*phaABC* Δ*mvaB* Δ*hbdH*, XW15) as a base, and performed the knockout of each gene from the previous prediction (*gltA* (citrate synthase), *aceA* (isocitrate lyase), and *gntZ* (6-phosphogluconate dehydrogenase, **Figure 3a**)). Since citrate synthase is the first enzyme connecting glycolysis and the TCA cycle, it plays an important role in central carbon and energy metabolism. In *P. putida* KT2440, citrate synthase derives 3-fold more carbon flux from acetyl-CoA to TCA cycle compared with *E. coli*^19^. Thus, the knockout of *gltA* may limit the flux out of acetyl-CoA, which is desirable to support the MVA pathway flux and isoprenol production. However, experimental results (**Figure 3b**) showed that the deletion of *gltA* (XW16) significantly compromised cell growth and eventually lowered isoprenol production (19 mg/L). Given that the clear detriment of cell growth, *gltA* was not continued with a further round of gene knockout. Adaptive laboratory evolution to increase growth or a less severe knockdown approach using CRISPRi^30^ are good candidate approaches to be attempted in the future. The knockout of *aceA* or *gntZ* on the XW15 strain, however, was still producing low levels of isoprenol: 12 mg/L for XW17 strain (Δ*phaABC* Δ*mvaB* Δ*hbdH* Δ*aceA*) and 13 mg/L for XW18 strain (Δ*phaABC* Δ*mvaB* Δ*hbdH* Δ*gntZ*), respectively (**Figure 3**). In the final round of gene knockouts, the XW17 strain was used to integrate the *gntZ* deletion to create the XW19 strain (Δ*phaABC* Δ*mvaB* Δ*hbdH* Δ*aceA* Δ*gntZ*) that contains all gene knockouts that do not affect growth. Interestingly, the inclusion of *gntZ* knockout significantly restored isoprenol production to 114 mg/L (**Figure 3b**). Although this isoprenol level was still lower than the two previous strains (XW12 and XW15), it reflects the synergy among multiple gene knockout targets suggested by the modeling predictions. To understand the different effects caused by Δ*aceA* and/or Δ*gntZ*, we compared the production profiles for XW17 to XW19 strains (**Supplementary Figure 1**). It was observed that the XW19 strain depleted glucose after 48 hours while the XW17 strain showed 6 g/L residual glucose in the medium. Correspondingly, the cell growth showed the opposite trends in these three strains, such that the XW19 strain showed the highest OD, which is 1.7-fold higher than that of the XW17 strain.

While the model requires knockout of all targets provided, we instead selected a subset of these targets as ranked by the frequency provided by several methods. The deletion of a subset of selected targets still showed significant improvement for isoprenol production even though the model does not predict growth-coupled isoprenol production with a subset. While a correlation analysis between isoprenol production and cell growth (OD_600_) only showed a weak positive correlation (P<0.05, R^2^=0.47), it was observed that the higher producers usually showed better cell growth (**Supplementary Figure 2**). This suggested that the increased isoprenol production is not at the expense of cell growth at the current isoprenol levels. However, not all strains showed improved isoprenol production and it might be necessary to delete additional genes to achieve a higher yield. For example, the rebound of isoprenol production by integrating both *aceA* and *gntZ* in one strain shows the importance of synergetic effects among different gene targets. We also note that the model predictions were made using a minimal medium, whereas experiments were performed using EZ rich defined medium due to better isoprenol production^19^. Thus, optimizing the production performance in a minimal medium could be a prerequisite to further demonstrate the effectiveness of our frequency-based prioritization of gene targets. Nonetheless, these results show that the deletion of prioritized gene targets can lead to significant improvement in isoprenol production and thus provide support for our computational approach.

In summary, following the genome-scale metabolic modeling recommendations, we constructed single and multiple knockout *P. putida* mutant strains. Among the engineered knockout strains, a triple knockout mutant strain (XW15, Δ*phaABC* Δ*mvaB* Δ*hbdH*) showed the highest isoprenol production (241 mg/L) from EZ rich medium supplemented with 2% glucose. This demonstrated the possibility of the host strain optimization driven by computational approaches to develop an excellent producing platform to achieve high titer, rate, and yield.

### Pathway Optimization for Improved Isoprenol Production

In parallel with the metabolic rewiring efforts above, we continued to optimize the pathway gene expression to improve isoprenol production in *P. putida* KT2440. We first expressed the IPP-bypass isoprenol biosynthetic pathway comprising *mvaE, mvaS, mk,* and *pmd_HKQ_* in different plasmid backbones (**Figure 4a**; strains IY781-IY784, **Supplementary Table 1**) under the control of a LacI repressor in the *P. putida* Δ*phaABC* strain (XW01). The highest titer of 80 mg/L isoprenol was achieved from strain IY782, carrying the isoprenol pathway in the plasmid with the RK2 replication origin (**Figure 4b**). Therefore, in the subsequent experiments, plasmid RK2 was used as the expression vector. Since we identified that the arabinose-inducible promoter (P_BAD_) is a stronger promoter than P_A1lacO-1_, (**Supplementary Figure 3**) we expressed the isoprenol biosynthetic pathway under the P_BAD_ promoter and replaced the LacI repressor with AraC, in an effort to improve isoprenol production. The absence of the LacI repressor resulted in the constitutive expression of MK and PMD_HKQ_ under a strong P_trc-1O_ promoter^31^. The resulting strain (IY721) produced up to 395 mg/L isoprenol in 48 hr (**Figure 4b**).

**Figure 4.**
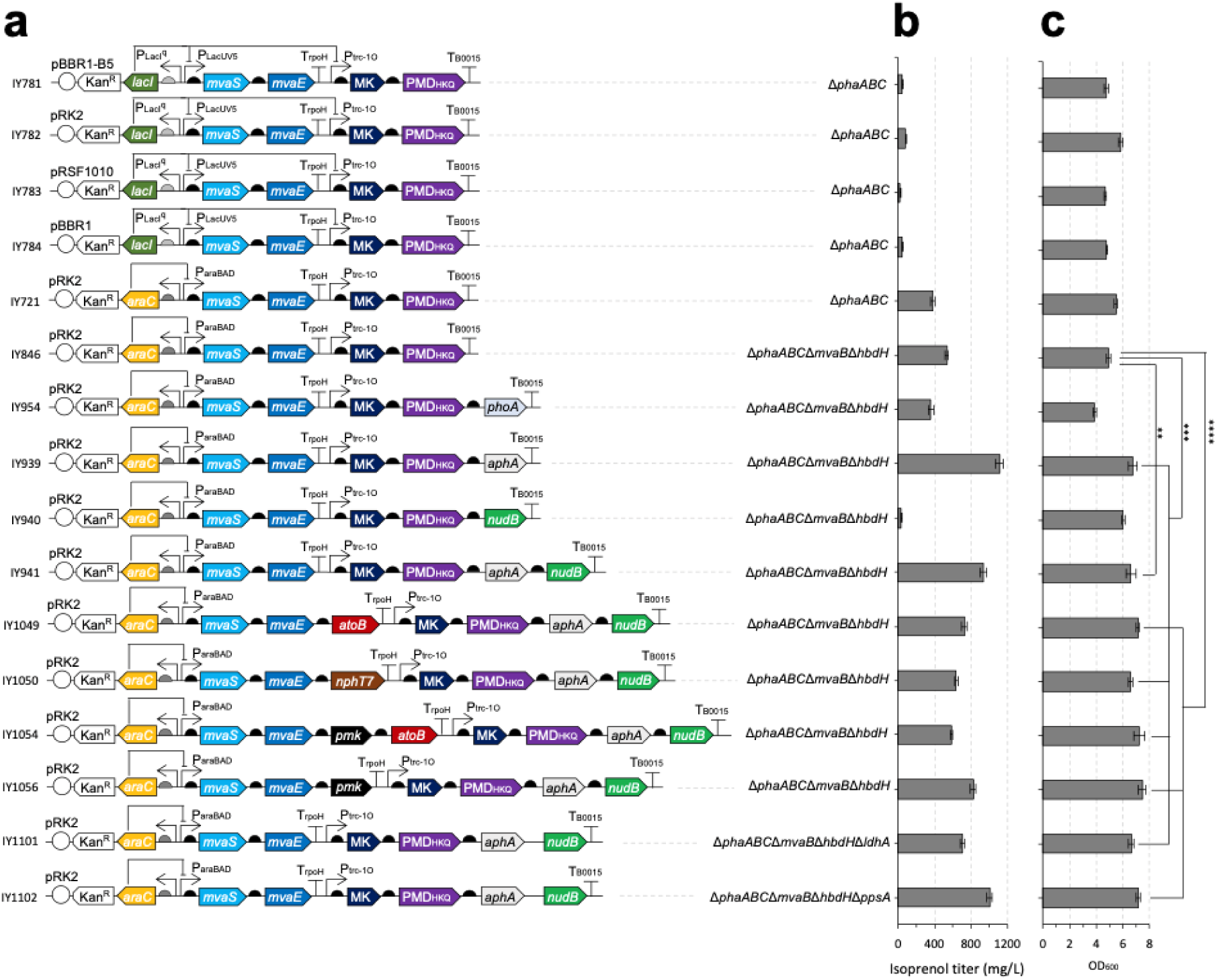
Pathway optimization in EZ Rich medium. **a** Schematic diagram of plasmids (Supplementary Table 2) used in this study. **b** Isoprenol titer obtained at 48 hr from strains transformed with plasmids shown in **a**. **c** Cell density (OD600) from strains shown in **b** measured at 48 h. Data were obtained from three biological replicates and error bars represent standard deviation. Asterisks indicate statistical significance by Student’s t-test (**, 0.001 < P < 0.01; *** 0.0001 < P < 0.001; **** P < 0.0001).

With a success on the pathway plasmid optimization, to further improve the production of isoprenol, we leveraged the knockout targets identified using the GSMM in previous sections and knocked out two genes: *mvaB* (encoding hydroxymethylglutaryl-CoA lyase) and *hbdH* (encoding 3-hydroxybutyrate dehydrogenase). The resulting strain (IY846) improved the isoprenol production by 1.3-fold, yielding approximately 536 mg/L isoprenol in 48 hr (**Figure 4b**).

Previous studies in *E. coli* have demonstrated that phosphatase overexpression can boost isoprenol production. Therefore, we further engineered the strains with coexpression of phosphatases. In our previous paper, we showed that NudB, a native phosphatase of *E. coli*, hydrolyzed IPP and DMAPP into their monophosphate forms, IP and DMAP, respectively, which are subsequently hydrolyzed to isoprenol by other phosphatases such as AphA, Agp, and YqaB^15^. Among these three phosphatases, AphA was found to best improve the isoprenol titer in *E. coli*^15^. A recent study for isoprenol production in *S. cerevisiae*, however, reported that co-expressing an *E. coli* alkaline phosphatase, PhoA, produced the highest isoprenol titer^11^. Therefore, we co-expressed NudB, PhoA, and AphA along with the isoprenol biosynthetic pathway and found that co-expressing AphA alone (strain IY939 with plasmid pIY670) produced the highest isoprenol titer of 1,111 mg/L in 48 hr (**Figure 4b**).

We also attempted to co-express AtoB, NphT7, and/or PMK, and knocked out *ldhA* and *ppsA* to increase the acetyl-CoA pool, and targeted proteomics confirmed the expression of all proteins (**Supplementary Figure 4**), but none of the strains generated higher isoprenol titer compared to strain IY939 (**Figure 4b**). Strains co-expressing AphA also accumulated higher biomass compared to those not expressing AphA (**Figure 4c**).

### Isoprenol Production Using Optimized Isoprenol Production Pathway and Predicted Metabolic Rewiring in Glucose Minimal Medium

Using the optimized isoprenol pathway, we continued to characterize the engineered strains carrying model-predicted gene knockouts. Defined rich medium such as EZ rich medium contains additional carbon sources such as amino acids and there could be complex regulation mechanisms such as carbon catabolite repression active^32–34^. The improved isoprenol production by the optimized pathway now enabled us to characterize the engineered strains in a minimal defined medium consistent with the condition used for predicting gene targets. We tested wild-type and 14 different knockout strains for growth and isoprenol production under M9 glucose minimal medium cultivation condition (**Figure 5**).

**Figure 5.**
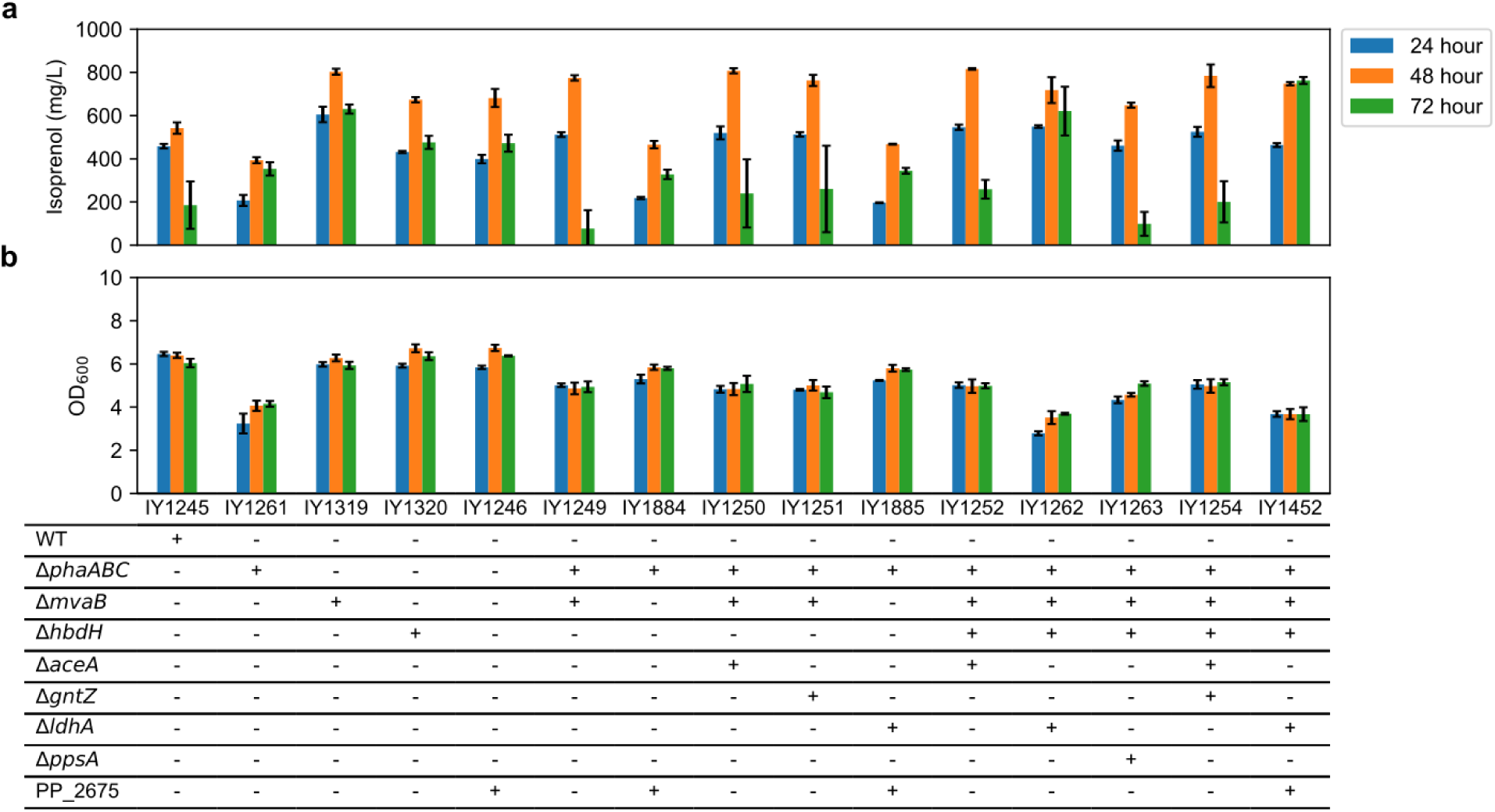
a. Isoprenol production and **b** growth in M9 glucose minimal medium using the optimized isoprenol production plasmid (pIY670, Supplementary Table 2) after double adaptation. Data were obtained from three biological replicates and error bars represent standard deviation.

We down selected the engineered strains harboring the heterologous, pathway optimized, IPP-bypass isoprenol production plasmid pIY670 based on their performance in EZ rich medium (discussed in the previous section) and produced isoprenol using M9 minimal medium containing 20 g/L glucose. Since we switched from EZ rich medium (several carbon sources) to minimal medium with glucose as the sole carbon source, we deleted an additional gene, PP_2675, to avoid the catabolism of isoprenol after glucose consumption as this gene was reported to be involved in isoprenol degradation^22^. We observed that the IY1452 strain (*ΔphaABC ΔmvaB ΔhbdH ΔldhA ΔPP_2675* with plasmid pIY670) had the best isoprenol titer (0.762 ± 0.016 g/L, **Figure 5**) that was over 4-fold higher than WT (IY1245) among the strains tested at 72 hr with 0.6-fold reduction in growth/biomass (WT, 6.04 ± 0.19; IY1452, 3.67 ± 0.31). The highest isoprenol titer observed was 0.816 g/L for the IY1252 strain at 48 h but that reduced by about 70% to 0.259 g/L at 72 hr. The maximum loss of isoprenol titers was observed for IY1249, a 90% reduction to 0.076 g/L at 72 hr from the titers observed at 48 hr time point (0.774 g/L). We observed that there was negligible reduction in isoprenol titers in the IY1452 strain at 72 hr unlike the other strains in M9 glucose minimal medium. Interestingly, the deletion of both *ldhA* and *PP_2675* was needed to improve the maximum isoprenol production titer while reducing the isoprenol degradation (**Supplementary Figure 5**).

Next we investigated the growth dynamics, carbon utilization, and isoprenol production profiles of the engineered strains by performing time-course profiling of the engineered strains (IY1245, IY1261, IY1249, IY939, IY1254, and IY1452) in M9 glucose minimal medium up to 72 hr. The growth rate and glucose consumption remained similar for all strains but there was a definite change in isoprenol production. The glucose was completely consumed by 42 hr in all the engineered strains (**Figure 6a**). The growth rate ranged between 0.22 hr^-1^ to 0.26 hr^-1^. There were changes in isoprenol production profiles for all of the strains. WT and IY1261 had similar isoprenol titers during early time points (glucose consumption phase) whereas strains IY1249, IY939, IY1254, and IY1452 had higher isoprenol production until ∼40 hr. Compared to the other *P. putida* strains, IY1452 sustains the isoprenol titers even after 40 hr when glucose is completely consumed. Data generated from 0 hr to 12 hr was used to estimate the rates for growth, glucose consumption, and isoprenol production (**Figure 6b**). IY1452 strain also had the best isoprenol production rate of 0.34 mmol/gCDW/hr which is a 4.69-fold improvement compared to WT strain harboring the production plasmid (IY1245) which had an isoprenol production rate of 0.07 mmol/gCDW/hr (**Figure 6b**).

**Figure 6.**
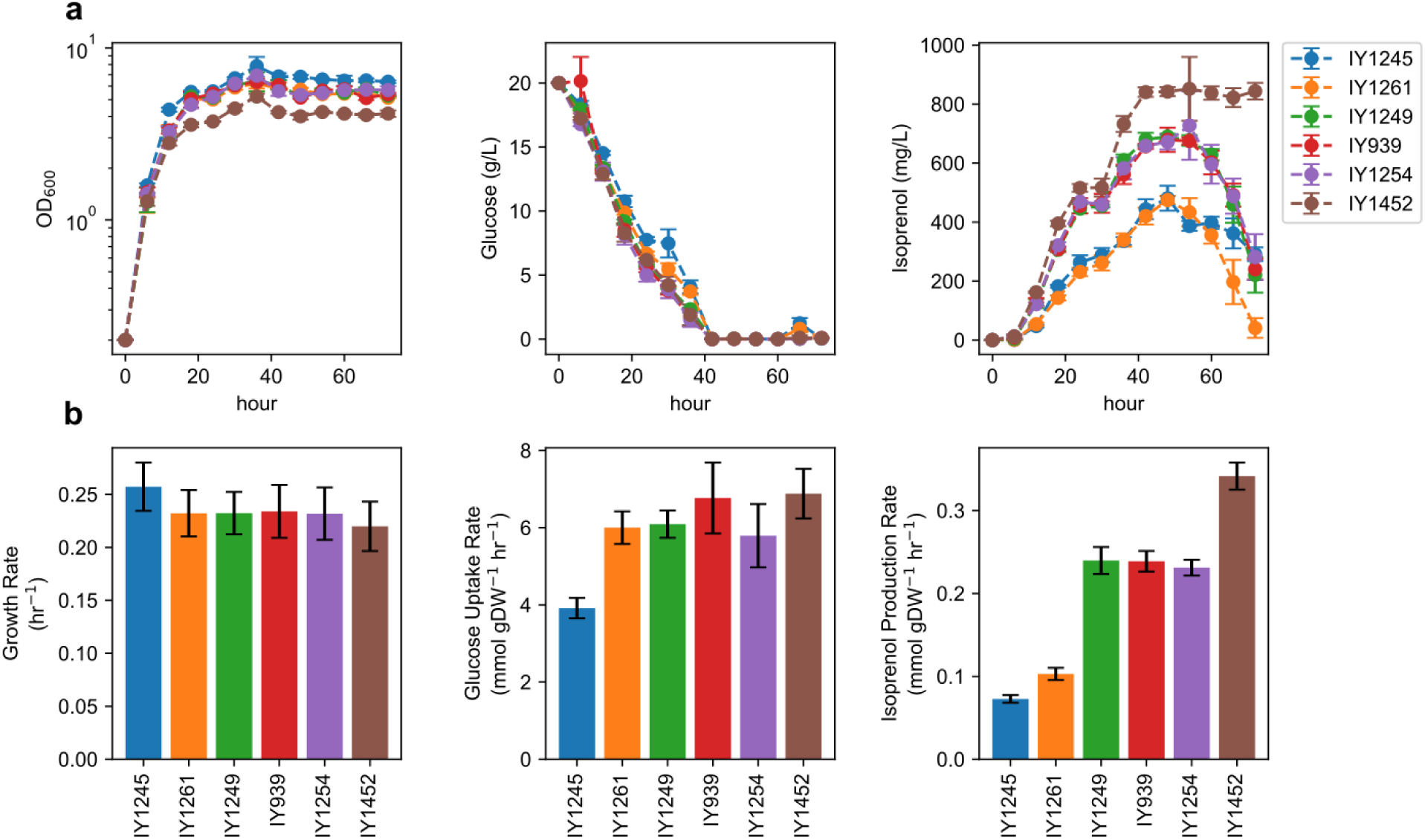
Time-course profiles of the engineered *P. putida* KT2440 strains in M9 glucose minimal medium. **a** Growth, glucose consumption, and isoprenol production. Data were obtained from three biological replicates and error bars represent standard deviation. **b** Specific rates estimated using the first three time points (0 hr, 6 hr and 12 hr) during exponential growth. Error bars represent the estimated standard error.

### Improved Isoprenol Producing Phenotype Observed for the IY1452 Strain

The phenotypic data that included experimentally measured glucose consumption, biomass formation, and isoprenol production rates (**Figure 6b**) were used to investigate the metabolic changes in the engineered strains using a GSMM. For this purpose, context-specific models were created for wild-type and engineered strains using constraints based on the gene knockouts as well as the phenotypic data. To compare the metabolic changes between the different engineered strains and WT, through flux redistribution, we performed flux variability analysis for each context-specific GSMM. We used flux variability analysis to assess the metabolic flux span for each of the reactions in a GSMM during optimal growth under the defined constraints and normalized it by the glucose uptake rate to be able to compare the variability across different GSMMs. The flux span corresponds to the flexibility/rigidity of each reaction during maximal growth and isoprenol production based on experimental constraints. A narrow flux span means a rigid flux due to either a causal or correlational effect of the strain engineering for improved isoprenol production. A narrow flux points towards a very constrained flux for the given production level, and hence an interesting candidate for future overexpression targets. A narrow flux may also be an effect of the engineering to reduce carbon flux redirection away from isoprenol production pathway. On the contrary, a wider flux span reflects that these reactions have numerous permissible carbon flux values during the current growth and observed isoprenol production. These are interesting future candidates for deletion for improvement of isoprenol production in *P. putida*. Selected reactions involving the precursor metabolites with respect to central metabolism and their flux spans are shown in **Figure 7**. The flux variability analysis (FVA) of the context-specific GSMM shows that the second reaction in the heterologous IPP-bypass isoprenol production pathway, HMGCOAS (*mvaS*, Reaction 11 in **Figure 7**), has a narrow flux span in the strains with additional gene deletions on top of *ΔphaABC,* i.e. strains that have the PP_3540/*mvaB* gene deletion. Given the constraints and assumptions in the GSMM, we can imagine two possible reasons for this narrow flux span in the *ΔmvaB* strains compared to the wider flux span in WT and *ΔphaABC* strains. It could be a result of high flux in competing pathways that redirect carbon flux away from the metabolic pathways of interest to maintain similar growth rates as WT and *ΔphaABC* strains (**Figure 6b**, 0.22 hr^-1^ versus 0.26 hr^-1^) given the limited resources available in M9 glucose minimal medium. Alternatively, the reduction in flux span in strains with *mvaB* deletion is perhaps due to the elimination of a critical degree of freedom, truly constraining the flux profile. Firstly using FVA, we observed a high flux in the HMG-CoA consuming reaction, in case of WT and *ΔphaABC* strains (Reaction 12 in **Figure 7**), a competing reaction that redirects carbon flux away from the heterologous IPP-bypass pathway. This high allowable flux was reduced by the PP_3540/*mvaB* deletion. Secondly, based on FVA, we observed there was a high flux span in BDH (*hbdH*, Reaction 13 in **Figure 7**), in the direction diverting flux towards butanoate metabolism, away from the heterologous IPP-bypass pathway. This was resolved by the PP_3073/*hbdH* deletion on top of the *mvaB* deletion. In the GSMM, ACACT1r (Reaction 10 in **Figure 7**) was assumed to have flux only in the direction of IPP-bypass due to additional pIY670 plasmid-borne *mvaE* activity. Zero flux through ACALDtpp (Reaction 2 in **Figure 7**) represents reduction in secretion of byproducts such as acetaldehyde.

**Figure 7.**
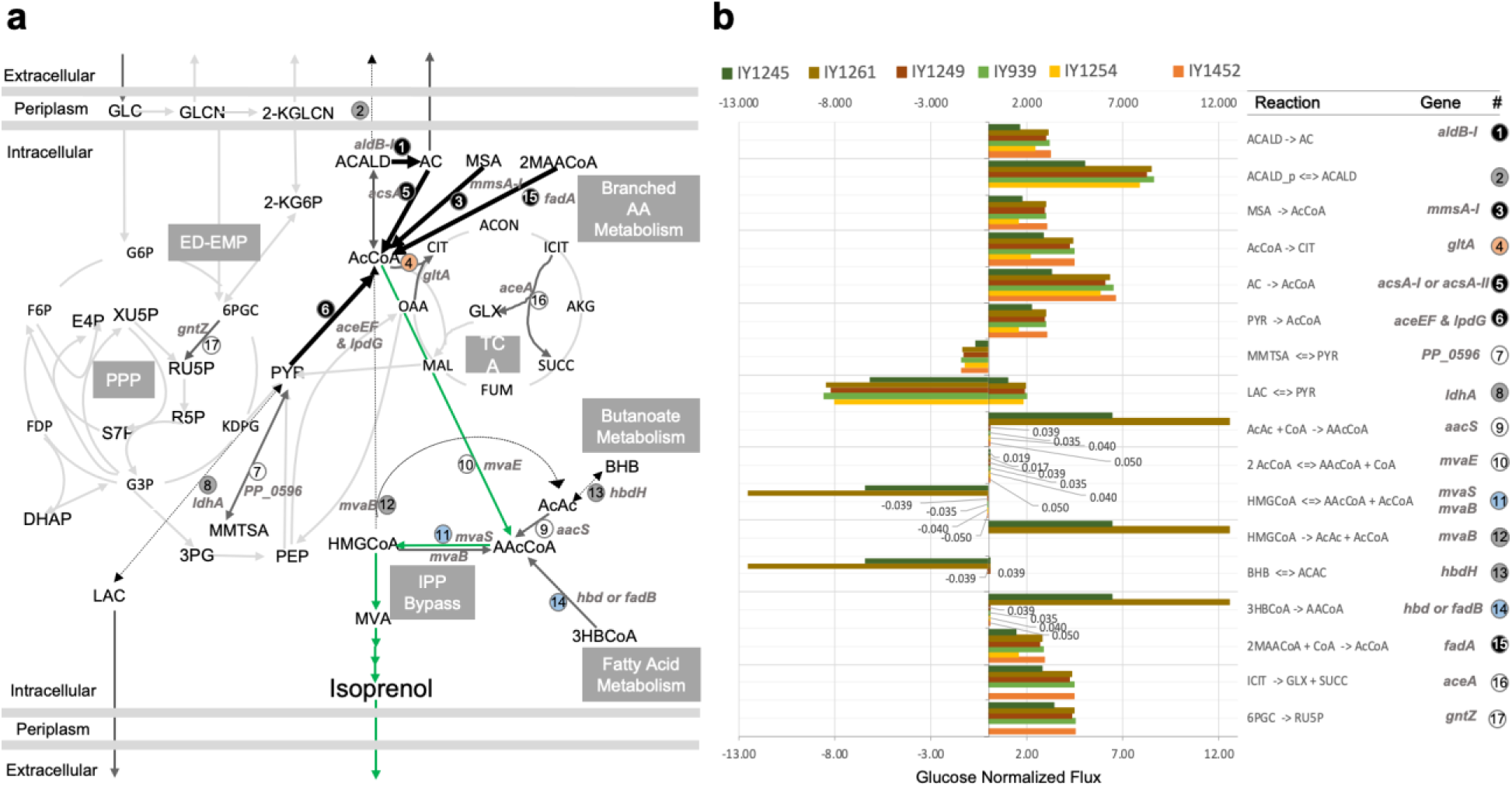
Flux variability analysis for the 6 different *P. putida* genotypes/strains with heterologous isoprenol production pathway (pIY670). **a** The central metabolic map is shown with reactions of interest highlighted and the heterologous isoprenol production pathway in green. **b** Flux span normalized to glucose uptake rate for selected reactions.

Although there were more than 50% blocked reactions (i.e., showing zero flux) in the *P. putida* KT2440 GSMM under glucose minimal medium conditions, 45% of the GSMM (1304 reactions) carried a substantial flux (**Supplementary Figure 6**). After constraining the model using our experimental data, the best performing strain IY1452 showed substantial fold changes in fluxes although the directionality of the reactions was similar to WT (**Supplementary Figure 7**). Of the 1,304 reactions in the GSMMs carrying a substantial flux, 964 reactions had an increase in the flux span by more than 1.5-fold compared to WT. These reactions with an increased carbon flow spread across central carbon metabolism (Glycolysis, Pentose Phosphate Pathway, Oxidative Phosphorylation, and TCA), fatty acid biosynthesis, most of the amino acid metabolism, nucleotide metabolism, glycerophospholipid metabolism, peptidoglycan biosynthesis, aromatic compounds degradation, heavy metal tolerance, and solvent extrusion transport metabolic subsystems. There were 225 reactions with a reduced flux span ranging from 1% to 80% of WT flux magnitudes. 69 of these reactions had 0.5-fold or lower flux. They mainly belong to alcohol degradation, butanoate metabolism, cell envelope lipopolysaccharide biosynthesis subsystems, and also the HMG-CoA synthase reaction of the IPP-bypass pathway. Additionally, there were about 156 reactions that had a flux span reduction between 0.5-fold to 0.8-fold. These were shared across non-unique subsystems including alanine and aspartate metabolism, branched amino acid metabolism, fatty acid metabolism, and PHA metabolism. Glyoxylate shunt through the ICL reaction and flux towards Pentose Phosphate Pathway through the GND reaction is substantial through all models except for IY1254 (*ΔphaABCΔmvaBΔhbdHΔaceAΔgntZ*), that had zero flux.

In summary, when compared to WT, the best performing strain IY1452 showed increased flux through desirable reactions for more acetyl-CoA pools (**Figure 7b**, reactions 1, 3, 5, 6, and 15) and reduced to negligible flux through competing reactions that redirect carbon flux away from isoprenol production pathway (**Figure 7b**, reactions 2, 8, 12 and 13). A restricted flux span through the reactions 11 and 14 point towards future strain engineering to redirect fatty acid metabolism towards generating HMG-CoA pools for further improvement in isoprenol production. A high flux span through CS (Reaction 4 in **Figure 7**) in most of the engineered strains point towards revisiting *gltA,* as a target for downregulation, in the best performing engineered *P. putida* strain for further improvement in isoprenol production. Although *gltA* deletion showed significant growth defect, CS was the top target predicted by the Opt-based method in this study (**Table 1**) and it was also previously identified by kinetic modeling as a target for downregulation to increase acetyl-CoA availability in *P. putida*^30^.

### Isoprenol Production in Fed-batch Cultivation

To increase isoprenol titer with additional supply of carbon and nitrogen sources (glucose and ammonium chloride, respectively), fed-batch mode production was performed with four strains (IY1245 (control), IY1262, IY1452, and IY1485). After the batch phase with the modified M9 minimal medium containing 20 g/L of glucose and 1.06 g/L (or 20 mM) ammonium chloride, an additional 200 mL feeding solution including 80 g glucose and 1.06 g ammonium chloride was continuously fed for fed-batch fermentation (a total of 100 g/L glucose and 2.12 g/L ammonium chloride was added). As isoprenol evaporates rapidly due to airflow in the bioreactor ^9^, the exhaust line was directly connected to a bottle containing 1 L oleyl alcohol as extraction solvent to extract isoprenol from the off-gas.

In the control strain, the maximum cell growth and the isoprenol production were obtained at 72 hr, reaching an OD_600_ of 25.5 ± 0.7 and isoprenol titer of 0.5 ± 0.1 g/L, respectively (**Figure 8**). Initial 20 g/L glucose was depleted by 14 hr and the isoprenol production was revealed from the off-gas after 24 hr, but no isoprenol was detected from the culture supernatant extract by then (Data not shown). This suggests that isoprenol produced in the vessel was evaporated by airflow as previously reported in the *E. coli* study^9^. The IY1262 strain produced 2.3 ± 0.3 g/L of isoprenol and the OD_600_ reached 20.5 ± 3.0 at 96 hr (**Figure 8**). The growth rate of the engineered strain was slower than the wild-type strain, but the titer was significantly increased. The maximum growth and isoprenol production on the IY1452 strain were obtained, reaching an OD_600_ of 21.0 ± 3.4 at 72 hr and a titer of 3.5 ± 0.3 g/L at 96 hr (**Figure 8**).

**Figure 8.**
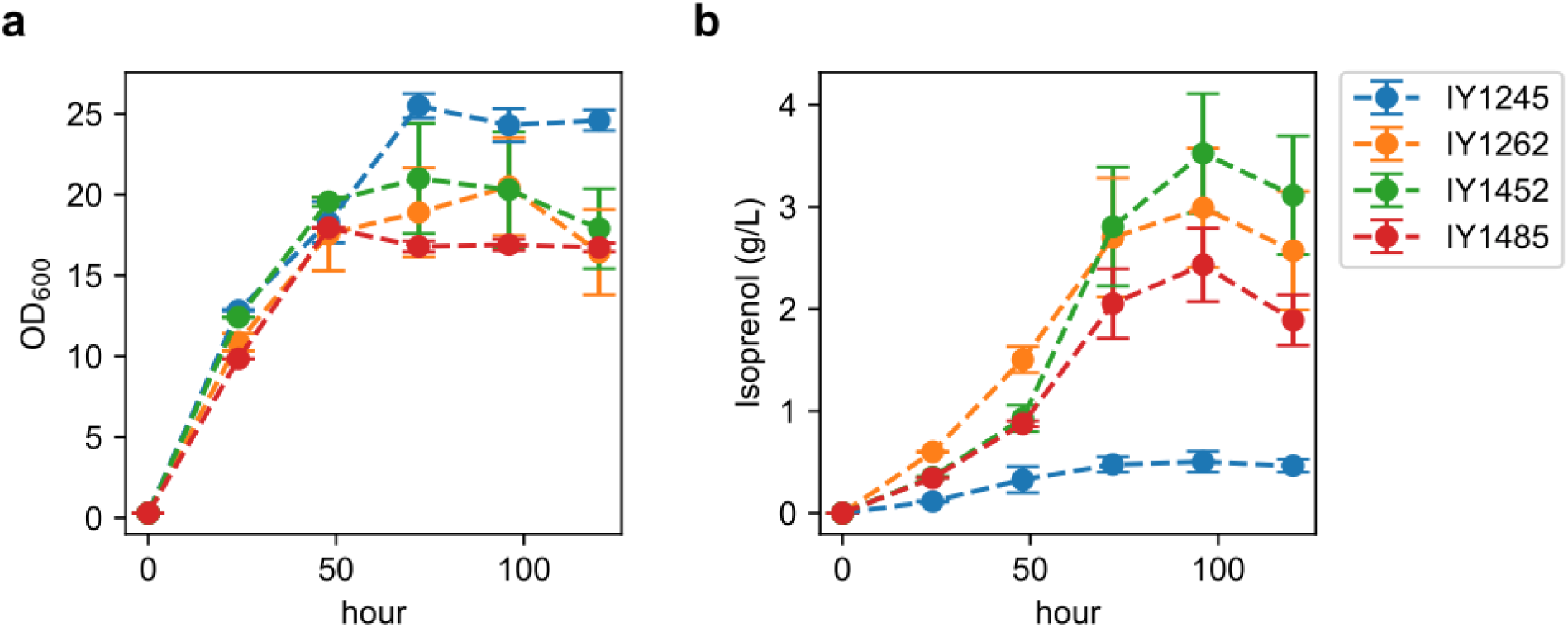
Growth and isoprenol production in fed-batch mode. **a** optical density (OD600) and **b** production of isoprenol by IY1245 (wild type), IY1262 (*ΔphaABCΔmvaBΔhbdHΔldhA*), IY1452 (*ΔphaABCΔmvaBΔhbdHΔldhA*ΔPP_2675), and IY1485 (*ΔphaABCΔmvaBΔhbdHΔldhA*ΔPP_2675*ΔgacA*). The fed-batch productions were performed in the 2 L bioreactors including M9 defined medium at triplicate. Error bars represent standard deviation.

Aeration is required during *P. putida* fed-batch cultivation, but it leads to strong foaming, which is technically difficult to handle even with conventional antifoams. Furthermore, the excessive foam formation hinders the use of standard cultivation protocol^35, 36^. To reduce foaming during the fed-batch cultivation, the *gacA* gene was deleted on the 4 genes knockout strain as previously reported^37^. The resulting IY1485 strain reached the OD_600_ of 17.9 ± 0.2 at 48 hr and produced 2.4 ± 0.3 g/L of isoprenol at 96 hr in fed-batch mode. Even though the *gacA* gene knockout resulted in a significant reduction of foaming, it also resulted in slower growth and lower isoprenol production than the other mutants. While most isoprenol production *P. putida* strain produced isoprenol at 20 % oxygen control, no isoprenol was produced at 30 % oxygen control (Data not shown), and this suggests that oxygen level is also very important for isoprenol production in *P. putida*.

### Isoprenol production using biomass hydrolysate

The use of lignocellulosic biomass for the production of biofuels and bioproducts is of increasing interest^38^ and *P. putida* is widely recognized for this purpose ^39, 40^. We tested whether biomass hydrolysates can be converted into isoprenol by our engineered isoprenol producing *P. putida* strain.

We evaluated the production of isoprenol using a modified M9 minimal medium supplemented with glucose or sorghum hydrolysate as the carbon source. The IY1452 strain was used to produce isoprenol using sorghum hydrolysate. The highest isoprenol titer from this strain was 0.841 g/L, which occurred at 72 hr in a modified M9 minimal medium supplemented with 20 g/L of glucose as a sole carbon source (**Figure 9a**). Isoprenol production using sorghum hydrolysate was lower than that with pure glucose, and the titers were 0.409 g/L and 0.432 g/L at 72 hr with 5% and 10% sorghum hydrolysate (matched the final glucose concentration of 20 g/L by adding pure glucose stock), respectively (**Figure 9a**). However, it is noteworthy that the growth rates of cultures with sorghum hydrolysate were higher than the growth rate with pure glucose (**Figure 9b**). Despite its lower isoprenol yield, the culture with sorghum hydrolysate showed promise as an alternative production medium with a higher growth rate than the culture with glucose as the sole carbon source.

**Figure 9.**
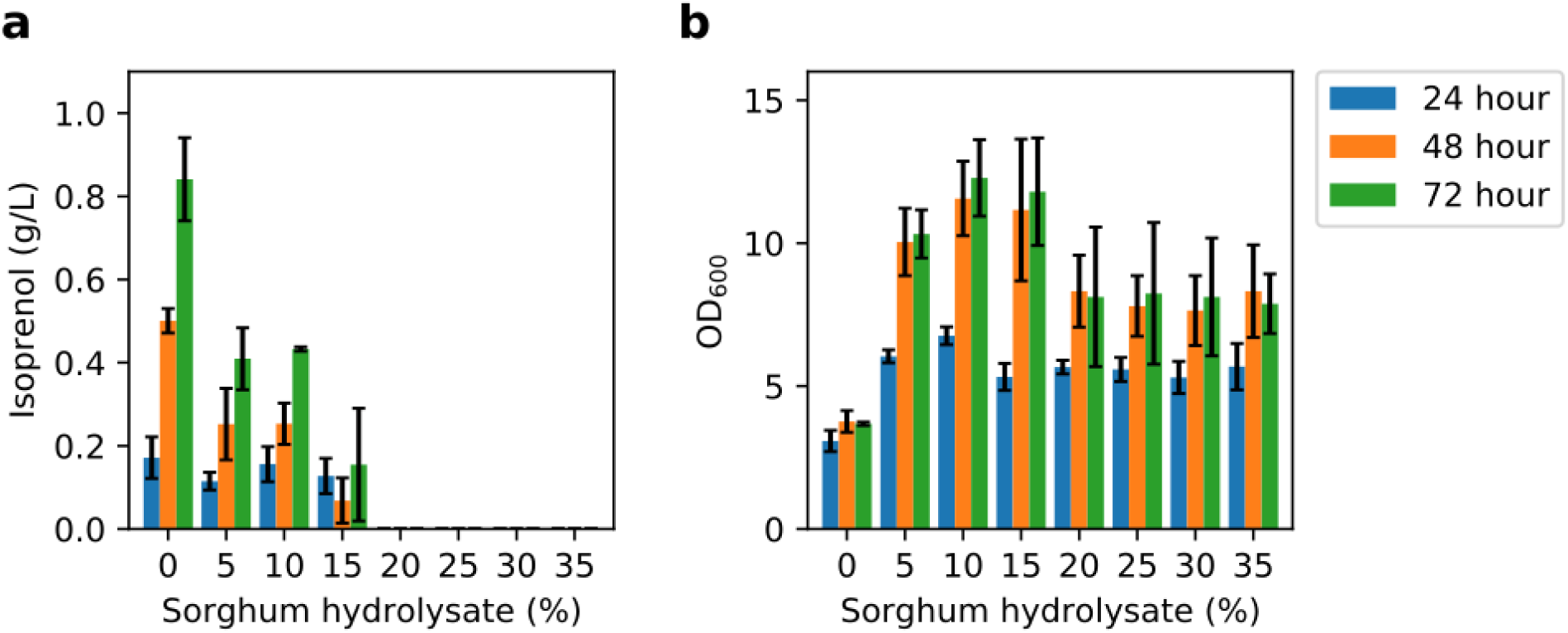
Isoprenol production and growth using hydrolysate. **a** production of isoprenol by IY1452 and **b** optical density (OD600). The productions of isoprenol and growth were evaluated in the culture tubes including 5 mL modified M9 minimal medium with varying concentrations of hydrolysate at triplicate. Error bars represent standard deviation.

Our GSMM-based computational strain design predictions were based on glucose as the sole carbon source under minimal medium cultivation conditions. Sorghum-based hydrolysate can be composed of different carbon sources based on the pretreatment methods^10, 41, 42^. It is reported that Sorghum-based ionic liquid ([Ch][Lys]) pretreated hydrolysate consists of glucose, xylose, acetate, and several aromatic compounds^10^. *P. putida* KT2440 lacks the capability to utilize xylose natively but has been reportedly engineered for xylose utilization^43, 44^. We observed improvement in growth across all tested fractions of hydrolysate but the isoprenol titers decreased with increasing fraction of hydrolysate in the medium when compared to glucose as the sole carbon source. This can be attributed to the presence of multiple carbon re-routing metabolic pathways towards growth versus limited bioconversion routes towards isoprenol production via the IPP-bypass pathway. For addressing the multi-carbon co-utilization of sorghum-based hydrolysate, medium constraints mimicking the hydrolysate carbon source composition for GSMM-based predictions hold promise in improving isoprenol production in the near future. Further, adaptive laboratory evolution (ALE)-based tolerization^41^ and other state-of-the-art strain engineering techniques^44, 45^ can also help improve isoprenol further to reach higher titers, rates, and yields.

## Conclusion

Global warming is an imminent crisis that threatens our survival and the efforts to reduce or slow down this threat are meaningful. In the aviation sector, which is hard to electrify, sustainable aviation fuels (SAFs) offer an effective near-term solution to global warming. In this study, we have reported our efforts to develop microbial hosts that can produce a SAF precursor utilizing broad carbon sources derived from biomass. We engineered *P. putida* strains via a combination of computational modeling and experimental execution of the prediction. Two computational modeling approaches, each of which could predict a handful of gene knockout targets, are evaluated and used to select and prioritize the intervention targets out of a much larger number of targets. This down-selection process significantly decreases the number of knock-out experiments to cover all predictions. However, we also observed that not all of these predictions contributed to titer improvement, and some combinations of knockouts were detrimental to growth and titer improvement. Pathway optimization had a significant impact on titer improvement, and the synergy of model-guided gene knockouts and pathway optimization led to the highest titer of isoprenol production in *P. putida* at 1.1 g/L which is over 10-fold improvement from the starting strain. Fed-batch cultivation improved the titer to 3.5 g/L, but for industrial application, it still requires more process optimization including downstream recovery of the volatile product.

As it is not a trivial task to knock out multiple genes in *P. putida* and the knock-outs frequently result in growth retardation, the gene knock-down could be an alternative to gene knock-out to screen multiple combinations of target genes. Application of CRISPR interference and building an automated process may accelerate rapid strain engineering to improve isoprenol production and enable a supply of the SAF to reduce CO_2_ emission in aviation.

## Methods

### Computation of constrained minimal cut sets (cMCS) and elementary modes

*Pseudomonas putida* KT2440 genome scale metabolic model (GSMM) iJN1462^24^ was used. Aerobic conditions with glucose as the sole carbon source were used to model growth parameters. The ATP maintenance demand and glucose uptake were 0.97 mmol ATP/gDW/h and 6.3 mmol glucose/gDW/h, respectively. Constrained minimal cut sets (cMCS) were calculated using the MCS algorithm^26^ available as part of CellNetAnalyzer (version 2020.2)^46^. Excretion of byproducts was initially set to zero, except for the reported overflow metabolites for secreted products specific to *P. putida* (gluconate, 2-ketogluconate, 3-oxoadipate, catechol, lactate, methanol, CO2, and acetate). We calculated the maximum theoretical yields (MTY) for isoprenol using glucose as the carbon source and the heterologous IPP bypass pathway (0.72 mol/mol of glucose). Additional inputs including minimum demanded product yield (10% to 85% of MTY) and maximum demanded biomass yield at 10 to 25% of maximum biomass yield were also specified in order to constrain the desired design space. The maximum size of MCS was kept at the default (i.e. 50 metabolic reactions). Knockouts of export reactions and spontaneous reactions were not allowed. With the specifications used herein, each calculated knockout strategy (cMCS) demands production of isoprenol even when cells do not grow. All cMCS calculations were done using API functions of CellNetAnalyzer^46^ on MATLAB 2017b platform using CPLEX 12.8 as the MILP solver. The different runs, respective number of cut sets and number of targeted reactions to satisfy coupling constraints are included in <u>Supplementary File 3</u>.

For elementary modes computation, a small model representing the central carbon metabolism of *Pseudomonas putida* KT2440 and the heterologous IPP bypass isoprenol production pathway was used to calculate elementary modes by *efmtool*^25^.

### Prediction of gene targets using opt-based methods

The iJN1462 metabolic model was also used for OptKnock and OptForce. The model was first modified to fix mass or charge unbalanced reaction, remove duplicate reactions involving lipoamide dehydrogenase, remove the reactions catalyzed by genes on the TOL plasmid pWW0, remove the PPCK reaction by a pseudogene phosphoenolpyruvate carboxykinase PP_0253, and update the gene-protein-reaction association for the OAADC reaction from 2-dehydro-3-deoxy-phosphogluconate aldolase PP_1024 to oxaloacetate decarboxylase PP_1389. The modified model was augmented with the IPP-bypass pathway for isoprenol production (<u>Supplementary File 1</u>).

For OptKnock, the model was preprocessed to remove blocked reactions and metabolites and identify the reactions predicted to be essential for growth on glucose as a sole carbon source. The predicted essential reactions, spontaneous reactions, boundary reactions, and periplasmic transport reactions without associated genes were excluded from knockout targets. Several additional reactions were manually excluded from knockout targets to avoid undesired predictions (ATPM, CAT, CYO1_KT, CYTBO3_4pp, CYTCAA3pp, NADH16pp, NAt3_1p5pp, PItex, and PPK).

The OptKnock problem was constructed using cobrapy^47^ and solved using CPLEX 12.8. Several iterations of OptKnock were run to identify a large number of knockout strategies using the solution pool and integer cuts. Another set of OptKnock solutions were obtained using a further constrained model where the secretion of other byproducts was blocked except for gluconate, 2-ketogluconate, and acetate assuming no significant byproduct formation (<u>Supplementary File 4</u>).

For OptForce, the model was also preprocessed to remove blocked reactions in glucose minimal media condition. The flux ranges for wild type were obtained by running flux variability analysis with constraints on glucose uptake, gluconate secretion, glucose dehydrogenase, gluconokinase, phosphogluconate dehydratase, pyruvate dehydrogenase, and citrate synthase taken from a previous study^48^. For isoprenol overproduction, we used 50 % of the theoretical maximum production as a pre-specified target to identify designs. All possible first and second-order necessary flux changes for overproduction were first identified and then used to identify the minimum set of interventions including flux increase, decrease, or knockouts. Several iterations of OptForce were run to identify a large number of designs by adding integer cuts using CPLEX 12.8 (<u>Supplementary File 4</u>).

### Context specific models and flux variability analysis

For flux variability analysis, first context-specific models were generated using constraints derived from experimental data to create six different *P. putida* GSMMs to represent the 6 different *P. putida* strains engineered for isoprenol production in this study. Constraints for glucose uptake rate, isoprenol production rate as well as growth rate were used. Next we performed flux variability analysis using *fluxvariabilityanalysis()* function in the COBRA Toolbox^49^ on the MATLAB 2017b platform.

### Strains and plasmid construction

All strains and plasmids used in this study are listed in **Supplementary Table 2**. Strains and plasmids along with their associated information have been deposited in the public domain of the JBEI Registry (https://public-registry.jbei.org) and are available from the authors upon request. Gene knockout of *P. putida* was performed based on the homologous recombination followed by a suicide gene (*sacB*) counter-selection as described^50^. The genotypes of gene-knockout mutants were confirmed by colony PCR using specific primers, followed by DNA sequencing (GENEWIZ, South San Francisco, CA, USA).

### Isoprenol production in *P. putida*

*P. putida* KT2440 strains bearing isoprenol pathway plasmids (**Supplementary Table 2**) were used for isoprenol production. Starter cultures of all production strains were prepared by growing single colonies in LB medium containing 50 µg/mL kanamycin at 30 °C with 200 rpm shaking overnight. The starter cultures were diluted in 5 mL EZ rich defined medium (Teknova, CA, USA) containing 20 g/L glucose (2%, w/v), 25 µg/mL kanamycin in 50-mL test tubes, and 0.5 mM IPTG or Arabinose (2%) was added to induce protein expression with OD_600_ at 0.4 – 0.6. The *P. putida* cultures were incubated in rotary shakers (200 rpm) at 30°C for 48 hr.

For isoprenol production runs on M9 minimal medium (**Supplementary Table 3**) with 2% D-glucose, cryostocks were streaked to singles on LB agar plate with 50 µg/mL kanamycin at 30 °C. Single colonies were inoculated and grown overnight with shaking in 5 mL liquid LB medium supplemented with 50 µg/mL kanamycin at 30 °C and 200 rpm. Unless otherwise mentioned, all further cultivations were performed in the same format and conditions. 100 µL of these overnight LB grown cultures were back diluted into the minimal medium and grown for 24 hr. A second back dilution enabled complete adaptation in the minimal medium. For the production runs, the cells were inoculated at an initial OD_600_ of 0.2 and the isoprenol production pathway was induced with 2% Arabinose immediately after inoculation. Samples were collected every 6 hr until 72 hr in triplicates and analyzed for growth (OD_600_), isoprenol, residual glucose and organic acids.

The quantification of isoprenol was conducted as described in Kim et al., 2021 ^11^. Briefly, 100 µL of ethyl acetate containing 1-butanol (30 mg/L) as the internal standard was added to 100 µL of liquid cultures. The mixture was vortexed at 3000 rpm for 15 min and subsequently centrifuged at 21,130 X g for 3 min to separate the ethyl acetate phase from the aqueous phase. 1 µL of the ethyl acetate layer was analyzed by gas chromatography-flame ionization detection (GC-FID, Thermo Focus GC) equipped with a DB-WAX column (15 m, 0.32 mm inner diameter, 0.25 µm film thickness, Agilent, USA). The GC oven was programmed as follows: 40 °C to 100 °C at 15 °C/min, 100 °C to 230 °C at 40 °C/min finally, held at 230 °C for 2 min. The inlet temperature was 200 °C. Serial dilutions of isoprenol were prepared to determine the quantification of isoprenol in the samples.

The residual glucose and organic acids were analyzed using high performance liquid chromatography (HPLC, Agilent, USA) equipped with a refractive index detector (RID) and an Aminex HPX-87X column (Bio-Rad, USA) with 4 mM sulfuric acid as the mobile phase in the isocratic mode. The following conditions were used: Mobile phase flow rate: 0.6 mL/min, column at 60 °C, RID at 35 °C. Serial dilutions of glucose and organic acids were used to determine the concentration of glucose and organic acids in the samples. Data analysis was carried out on the ChemStation software (Agilent Technologies).

### Targeted Proteomics Analysis of the Isoprenol Biosynthesis Pathway Proteins

Cell pellets of the engineered *P. putida* strains for isoprenol production were prepared for targeted proteomic analysis according to Chen et al^51^. Briefly, cells were resuspended in a solution with 80 μL of methanol and 20 μL of chloroform and thoroughly mixed by pipetting. Sixty microliters of water were subsequently added to the samples and mixed. Phase separation was induced with 5 minutes of centrifugation at 1000 x g. The methanol and water layers were removed, and then methanol (80 μL) was added to each well. The plate was centrifuged for 1 minute at 100 x g, and then the supernatant layers were decanted. The protein pellets were resuspended in a 100 mM ammonium bicarbonate buffer supplemented with 20% methanol, and the protein concentration was determined by the DC assay (Bio–Rad). proteins from each sample were reduced by addition of tris 2-(carboxyethyl) phosphine to 5 mM for 30 min at room temperature and followed by alkylation with iodoacetamide at 10 mM for 30 min at room temperature in the dark. Protein digestion with trypsin at 1 g/L concentration was accomplished with a 1:50 (w/w) trypsin/total protein ratio overnight. The multiple-reaction monitoring (MRM) assay was developed for relative quantification of isoprenol biosynthesis pathway proteins through a rapid method development workflow established previously^52^. Targeted proteomic analysis was performed on an Agilent 1290 UHPLC system coupled to an Agilent 6460 QqQ mass spectrometer according to an established protocol (<u>dx.doi.org/10.17504/protocols.io.bf9xjr7n</u>). Briefly, 20 g Peptides of each sample were separated on an Ascentis Express Peptide C18 column [2.7-mm particle size, 160-Å pore size, 5-cm length × 2.1-mm inside diameter (ID), coupled to a 5-mm × 2.1-mm ID guard column with the same particle and pore size, operating at 60 °C; Sigma-Aldrich] operating at a flow rate of 0.4 ml/min via the following gradient: initial conditions were 98% solvent A (0.1% formic acid), 2% solvent B (99.9% acetonitrile, 0.1% formic acid). Solvent B was increased to 5% over 1 min, and was then increased to 40% over 3.5 min. It was increased to 80% over 0.5 min and held for 2.5 min at a flow rate of 0.6 mL/min, followed by a linear ramp back down to 2% B at a flow rate of 0.4 mL/min over 0.5 min where it was held for 1 min to re-equilibrate the column to original conditions. The eluted peptides were ionized via an Agilent Jet Stream ESI source operating in positive ion mode. The MS raw data were acquired using Agilent MassHunter version B.08.02, and were analyzed by Skyline software version 21.20 (MacCoss Lab Software). The MRM method and data are available at Panoramaweb^53^ (https://panoramaweb.org/genome-scale-eng-SAF-p-putida.url) and at ProteomeXchange via identifier PXD039868.

### Isoprenol Production in Fed-batch Mode

For the isoprenol production in fed-batch mode, the strains were cultured in 5 mL LB medium with 50 µg/mL kanamycin at 30 ᵒC. For the adaptation, the cell culture was diluted 50-fold in the fresh 5 mL modified M9 minimal medium two times. Then the seed culture was inoculated in the 1 L baffled flask including 100 mL modified M9 minimal medium at 30 ᵒC and 200 rpm for 8 hr. The cell culture was inoculated at OD_600_ 0.3 in the 2 L bioreactor (Biostat B, Sartorius, Germany) including the 1 L modified M9 minimal medium, which contained 6.8 g/L Na_2_HPO_4_, 3.0 g/L KH_2_PO_4_, 0.5 g/L NaCl, 20 mM NH_4_Cl, 2 mM MgSO_4_, 0.1 mM CaCl_2_ and trace metal solution. The 1000X trace metal solution (TEKNOVA, USA) consisted of 50 mM FeCl_3_, 20 mM CaCl_2_, 10 mM MnCl_2_, 10 mM ZnSO_4_, 2 mM CoCl_2_, 2 mM CuCl_2_, 2 mM NiCl_2_, 2 mM Na_2_MoO_4_, 2 mM Na_2_O_3_Se, and 2 mM BH_3_O_3_. To produce isoprenol in the fed-batch fermentation, the dissolved oxygen (DO) and airflow were set to 20 % and 1 VVM (volume of air per volume of liquid per minute), respectively. The temperature was maintained 25 °C and pH was maintained at 6.5 by supplementation with 25% ammonia water. The isoprenol biosynthesis pathway was induced at OD_600_ 0.6 – 0.8 by 2 % arabinose. The antifoam B was added to the bioreactor when required. To feed the additional carbon and nitrogen sources, the feeding solution of 80 g glucose and 15 g ammonium chloride was continuously supplied using a Watson-Marlow DU520 peristaltic pump. The feeding flow rate was set to closely match the glucose consumption rate at the end of the batch phase. After the lag phase, the feeding flow rate was calculated following Korz’s equation and increased every hour for a total of 6 hr ^9,54^.

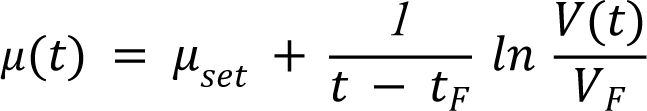

For the exponential feeding, the glucose in the media was measured consistently using the glucose meter (CVSHealth, USA) and high-performance liquid chromatography (HPLC), and feeding rate was set constant in order that the concentration of glucose was dropped below than 1 g/L in the medium. To extract the isoprenol from the off gas, the exhaust line was connected directly to a bottle including the 1 L oleyl alcohol as extraction solvent. For quantification of isoprenol, 10 μL of oleyl alcohol layer was added to 990 mL of ethyl acetate containing 1-butanol (30 mg/L) as internal standard.

### Isoprenol Production Using Biomass Hydrolysate

For the isoprenol production using biomass hydrolysate, the cholinium lysinate ionic liquid-pretreated sorghum hydrolysate was obtained from Joint BioEnergy Institute (JBEI) Deconstruction Division^55^ and was used as a carbon source for the engineered *P. putida* strain. The sorghum hydrolysate was adjusted at pH 6.5 by sodium hydroxide and supplemented with 10 X modified M9 salts at varying concentrations (0 %, 5 %, 10 %, 15 %, 20 %, 25 %, 30 %, and 35 % (v/v)). The glucose was added either as a sole carbon source or as co-substrate with sorghum hydrolysate and its concentration was adjusted to 20 g/L. Subsequently, we added 10 μL of 1 M MgSO_4_, 5 μL of 100 mM CaCl_2_, 2.5 μL of trace metal solution, and 50 μL of 1M NH_4_Cl to the modified M9 minimal medium. And the modified M9 minimal medium volume was adjusted to 5 mL. The strains were cultured in 5 mL LB medium with 50 µg/mL kanamycin at 30 ᵒC overnight. To adapt the strain, the cell culture was diluted 50-fold in the fresh 5 mL modified M9 minimal medium two times. Then the seed culture was inoculated in the culture tubes including 5 mL modified M9 minimal medium supplemented sorghum hydrolysate at 30 ᵒC and 200 rpm. The isoprenol biosynthesis pathway was induced at OD_600_ 0.6 – 0.8 by 2 % arabinose. The isoprenol extraction was carried out using the same protocols as described above.

## Supporting information

SI_File_1_iJN1462_modified

SI_File_2_PputidaCoreModel_IPPBypass

SI_File_3_cMCS_Solutions.xlsx

SI_File_4_Opt-based_Methods_Solutions.xlsx

SI_File_5_Combined_Scores

## Acknowledgements

This work was supported by the US Department of Energy, Office of Science, Office of Biological and Environmental Research, through Contract DE-AC0205CH11231 between Lawrence Berkeley National Laboratory and the US Department of Energy.

## Author Contributions

DB, IYS, XW and JoK contributed equally to the work. Conceptualization of the project: DB, JoK, AM and TSL. Development and implementation of computational methods: DB and JoK. Strain construction, molecular biology, analytical chemistry: ISY, XW, AS, RM. Proteomic analysis: YC, JWG, CJP. Interpreted results: DB, IYS, XW, JoK, TSL. Bioreactors and Scale-up: JiK. Writing-Original Draft: DB, ISY, XW, JoK, TSL. Writing-Review and Editing: DB, ISY, XW, JiK, HGM, AM, JoK, TSL. Funding Acquisition: TSL and AM. Supervision: TSL and AM. All authors contributed to and provided feedback on the manuscript. All authors read and approved the final manuscript.

## Supplementary Information

Supplementary Figures and Supplementary Tables are available in a separate pdf file (“Supplementary Information.pdf”). Additional Supplementary files are also available, and descriptions are listed below.

## Description of Additional Supplementary Files

Supplementary File 1. Genome-scale metabolic network model of *P. putida* KT2440 iJN1462 augmented with the heterologous MVA pathway for isoprenol production.

Supplementary File 2. Central metabolic network model of *P. putida* KT2440 augmented with a lumped reaction for the heterologous MVA pathway for isoprenol production.

Supplementary File 3. Computational designs for isoprenol production using elementary flux modes and constrained minimal cut sets.

Supplementary File 4. Computational designs for isoprenol production using OptKnock, and OptForce.

Supplementary File 5. Combined scores of the targets from EMA-based and Opt-based methods.

## Data Availability

All the experiments reported in this study are available through Experimental Data Depot (EDD) using the following links:

- https://edd.jbei.org/s/isoprenol-production-in-pseudomonas-kt2440-m9-medi/
- https://edd.jbei.org/s/isoprenol-production-in-pseudomonas-kt2440-ez-6943/
- https://edd.jbei.org/s/isoprenol-production-in-pseudomonas-kt2440-time-se/
- https://edd.jbei.org/s/isoprenol-degradation-in-pseudomonas-kt2440-time-s/
- https://edd.jbei.org/s/isoprenol-production-in-p-putida-try-project-fba1/
- https://edd.jbei.org/s/isoprenol-production-in-p-putida-using-biomass-hyd/

## Code Availability

All custom computer code used to generate the results reported in this manuscript are available via the Jupyter Notebooks and Supplementary Information provided along with this manuscript.

