## Supplementary Information - Figures and Tables for "Genome-scale and pathway engineering for the sustainable aviation fuel precursor isoprenol production in *Pseudomonas putida*"

### Supplementary Figures

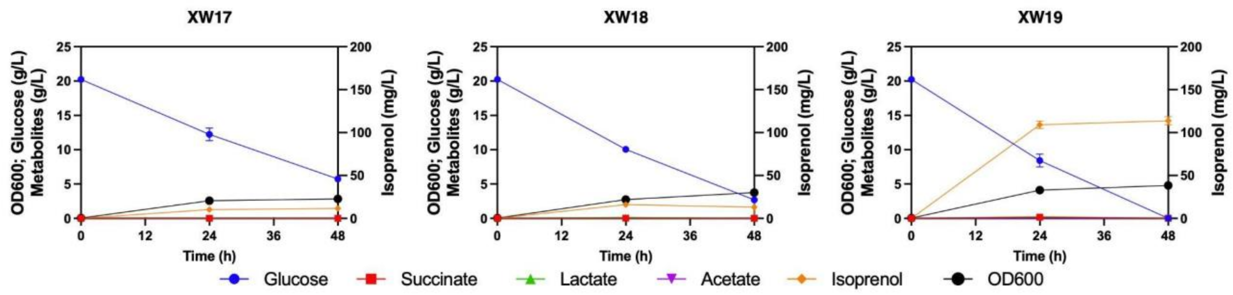


**Supplementary Figure 1:** Comparison of isoprenol production for multiple gene knockout strains. Time-course production of XW17 to XW19 strains in EZ rich media. Glucose, isoprenol, $OD_{600}$, and organic acids were measured. Data were obtained from three biological replicates and error bars represent standard deviation.

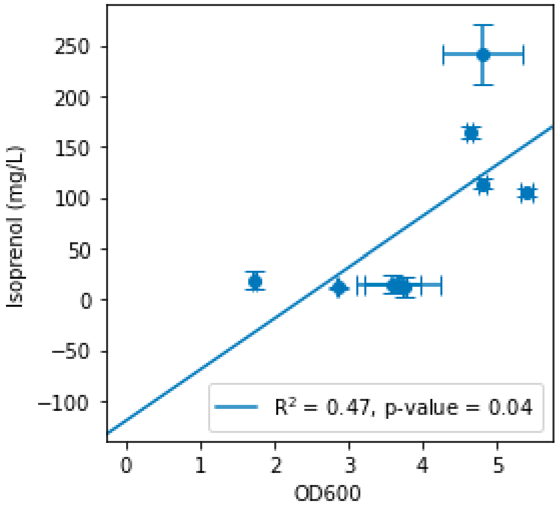


**Supplementary Figure 2:** Correlation between isoprenol production and cell growth of XW11 to XW19 strains in EZ rich media. Error bars represent standard deviation from three biological replicates.

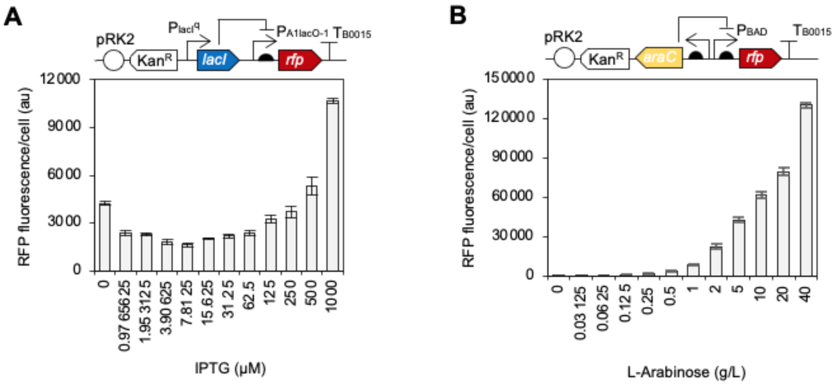


**Supplementary Figure 3:** Fluorescence level of red fluorescent protein expressed under (A) $P_{A1lacO-1}$ and (B) $P_{BAD}$ inducible promoter. Cultures were induced with different concentrations of IPTG or L-Arabinose, 2 hr after inoculation. Fluorescence level from at least 50000 cells was measured using a BD C6 Accuri flow cytometer with FL-4 detector at 24 hr. Error bars represent standard deviation of three biological replicates.

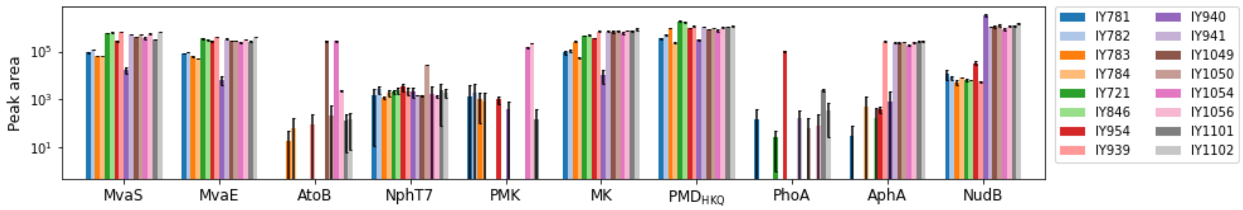


**Supplementary Figure 4:** Targeted proteomics of isoprenol production pathway. Data were obtained from three biological replicates and error bars represent standard deviation.

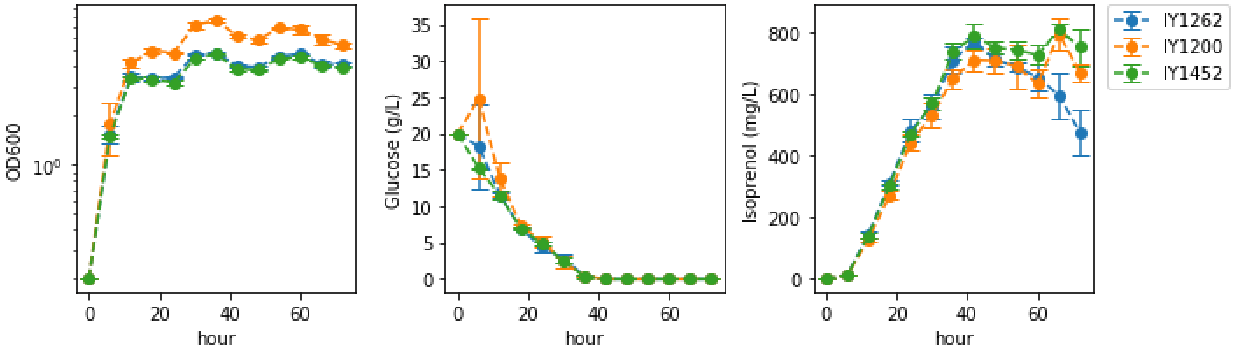


**Supplementary Figure 5:** Cell growth, glucose consumption, and isoprenol production/degradation from time-course experiment in M9 medium by IY1262 (*ΔphaABCΔmvaBΔhbdHΔldhA*), IY1200 (*ΔphaABCΔmvaBΔhbdHΔ*PP_2675), and IY1452 (*ΔphaABCΔmvaBΔhbdHΔldhA*ΔPP_2675). Data were obtained from three biological replicates and error bars represent standard deviation.

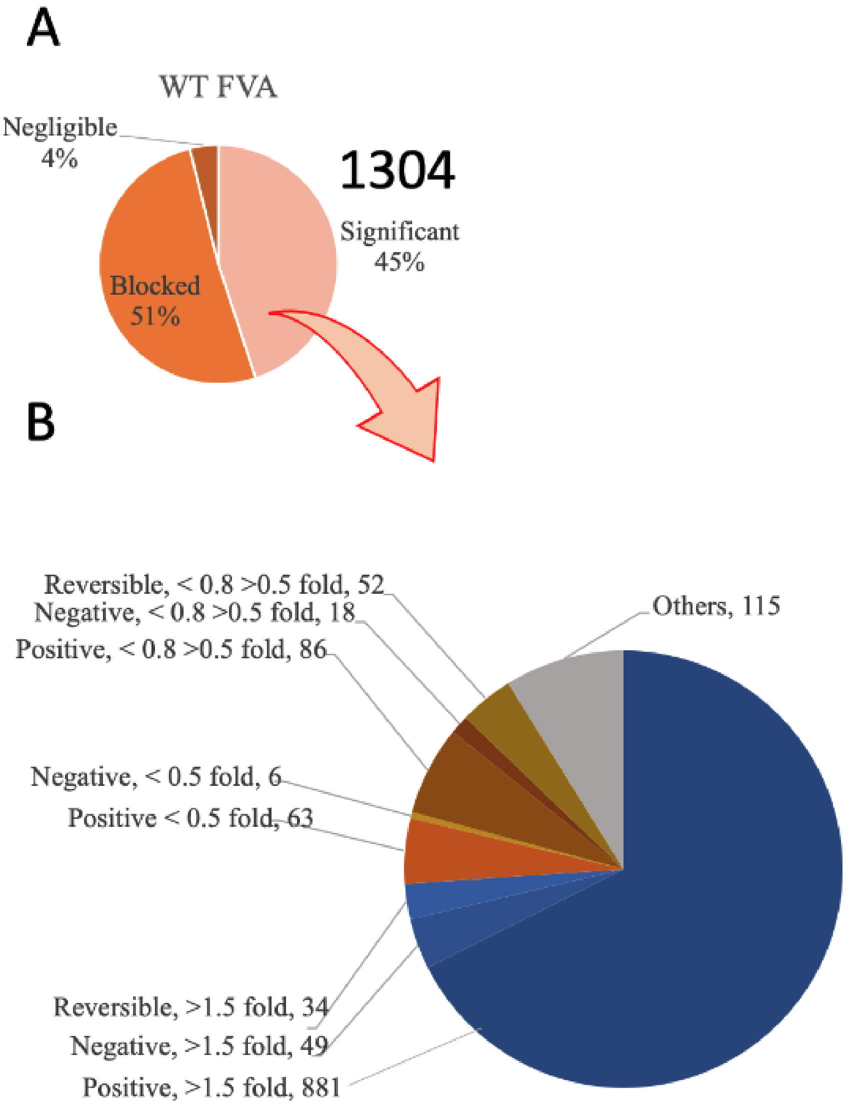


**Supplementary Figure 6:** Flux variability analysis (FVA) for *P. putida* KT2440 *ΔphaABCΔmvaBΔhbdHΔldhAΔ*2675 compared to WT flux span distribution normalized to glucose consumption.

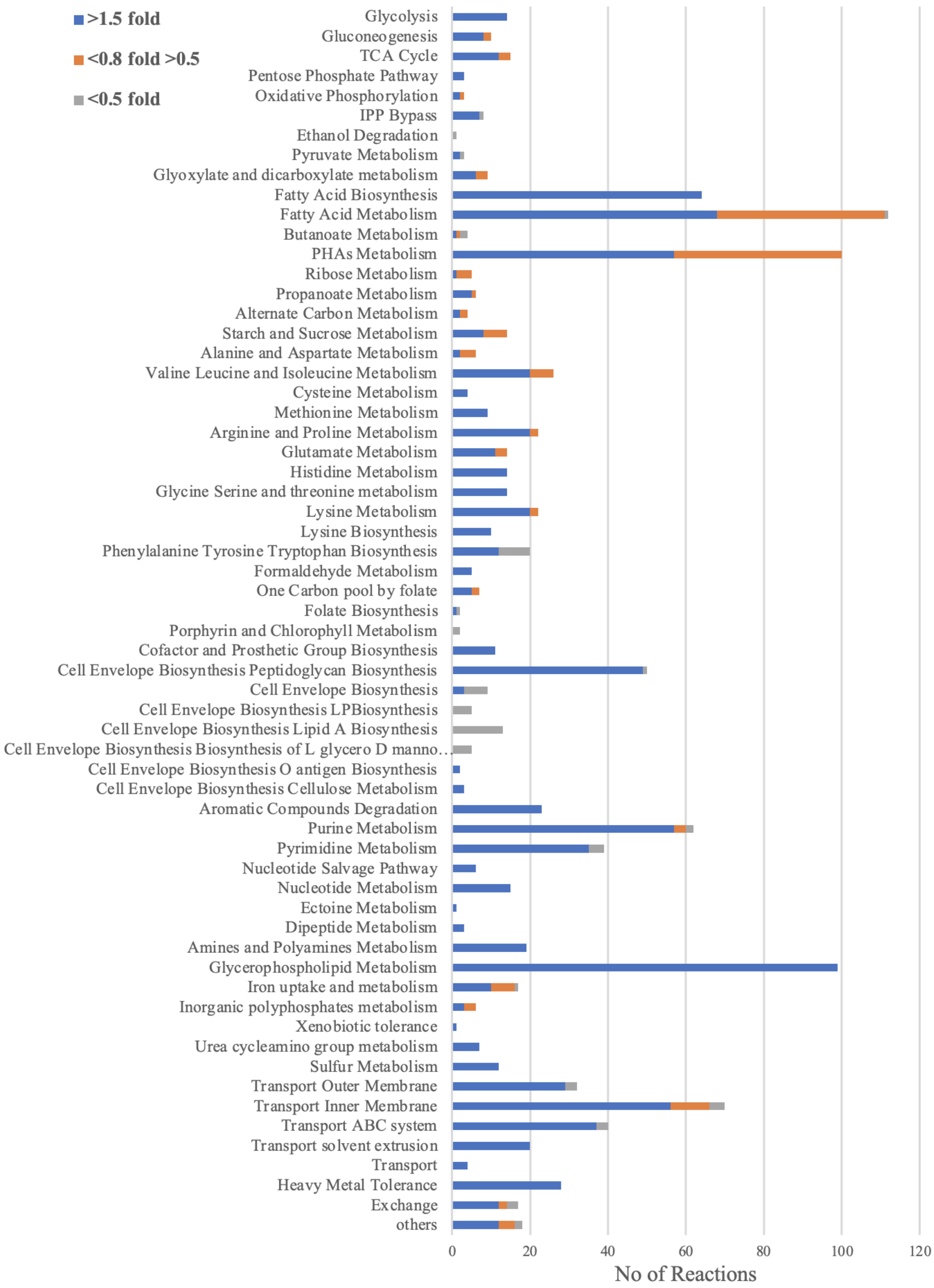


**Supplementary Figure 7:** Subsystem-wise distribution of reactions that had a substantial fold change in flux for *P. putida* KT2440 *ΔphaABCΔmvaBΔhbdHΔldhAΔ*2675 compared to WT.

### Supplementary Tables

**Supplementary Table 1.** Strains used in this study

| Strains | JBEI Registry | Description | Reference |
|---|---|---|---|
| XW01 | JPUB_019964 | *P. putida* KT2440 Δ*phaABC* | Wang et al. 2022 |
| XW02 | JPUB_019990 | *P. putida* KT2440 Δ*phaABC* Δ*mvaB* | This study |
| XW03 | JPUB_019992 | *P. putida* KT2440 Δ*phaABC* Δ*mvaB* Δ*aceA* | This study |
| XW04 | JPUB_019994 | *P. putida* KT2440 Δ*phaABC* Δ*mvaB* Δ*gntZ* | This study |
| XW05 | JPUB_019996 | *P. putida* KT2440 Δ*phaABC* Δ*mvaB* Δ*hbdH* | This study |
| XW06 | JPUB_019998 | *P. putida* KT2440 Δ*phaABC* Δ*mvaB* Δ*hbdH* Δ*gltA* | This study |
| XW07 | JPUB_020000 | *P. putida* KT2440 Δ*phaABC* Δ*mvaB* Δ*hbdH* Δ*aceA* | This study |
| XW08 | JPUB_020002 | *P. putida* KT2440 Δ*phaABC* Δ*mvaB* Δ*hbdH* Δ*gntZ* | This study |
| XW09 | JPUB_020004 | *P. putida* KT2440 Δ*phaABC* Δ*mvaB* Δ*hbdH* Δ*aceA* Δ*gntZ* | This study |
| XW11 | JPUB_019977 | *P. putida* KT2440 Δ*phaABC* with plasmid pXW1 | Wang et al. 2022 |
| XW12 | JPUB_019991 | *P. putida* KT2440 Δ*phaABC* Δ*mvaB* with plasmid pXW1 | This study |
| XW13 | JPUB_019993 | *P. putida* KT2440 Δ*phaABC* Δ*mvaB* Δ*aceA* with plasmid pXW1 | This study |
| XW14 | JPUB_019995 | *P. putida* KT2440 Δ*phaABC* Δ*mvaB* Δ*gntZ* with plasmid pXW1 | This study |
| XW15 | JPUB_019997 | *P. putida* KT2440 Δ*phaABC* Δ*mvaB* Δ*hbdH* with plasmid pXW1 | This study |
| XW16 | JPUB_019999 | *P. putida* KT2440 Δ*phaABC* Δ*mvaB* Δ*hbdH* Δ*gltA* with plasmid pXW1 | This study |

| | | | |
|---|---|---|---|
| XW17 | JPUB_020001 | *P. putida* KT2440 Δ*phaABC* Δ*mvaB* Δ*hbdH* Δ*aceA* with plasmid pXW1 | This study |
| XW18 | JPUB_020003 | *P. putida* KT2440 Δ*phaABC* Δ*mvaB* Δ*hbdH* Δ*gntZ* with plasmid pXW1 | This study |
| XW19 | JPUB_020005 | *P. putida* KT2440 Δ*phaABC* Δ*mvaB* Δ*hbdH* Δ*aceA* Δ*gntZ* with plasmid pXW1 | This study |
| IY721 | JBEI-233609 | *P. putida* KT2440 Δ*phaABC* with plasmid pIY554 | This study |
| IY781 | JBEI-233601 | *P. putida* KT2440 Δ*phaABC* with plasmid pIY602 | This study |
| IY782 | JBEI-233603 | *P. putida* KT2440 Δ*phaABC* with plasmid pIY603 | This study |
| IY783 | JBEI-233605 | *P. putida* KT2440 Δ*phaABC* with plasmid pIY604 | This study |
| IY784 | JBEI-233607 | *P. putida* KT2440 Δ*phaABC* with plasmid pIY605 | This study |
| IY846 | JBEI-233611 | *P. putida* KT2440 Δ*phaABC* Δ*mvaB* Δ*hbdH* with plasmid pIY554 | This study |
| IY939 | JBEI-233615 | *P. putida* KT2440 Δ*phaABC* Δ*mvaB* Δ*hbdH* with plasmid pIY670 | This study |
| IY940 | JBEI-233617 | *P. putida* KT2440 Δ*phaABC* Δ*mvaB* Δ*hbdH* with plasmid pIY671 | This study |
| IY941 | JBEI-233619 | *P. putida* KT2440 Δ*phaABC* Δ*mvaB* Δ*hbdH* with plasmid pIY672 | This study |
| IY954 | JBEI-233613 | *P. putida* KT2440 Δ*phaABC* Δ*mvaB* Δ*hbdH* with plasmid pIY697 | This study |
| IY1049 | JBEI-233621 | *P. putida* KT2440 Δ*phaABC* Δ*mvaB* Δ*hbdH* with plasmid pIY761 | This study |
| IY1050 | JBEI-233623 | *P. putida* KT2440 Δ*phaABC* Δ*mvaB* Δ*hbdH* with plasmid pIY762 | This study |

| | | | |
|---|---|---|---|
| IY1054 | JBEI-233625 | *P. putida* KT2440 Δ*phaABC* Δ*mvaB* Δ*hbdH* with plasmid pIY765 | This study |
| IY1056 | JBEI-233627 | *P. putida* KT2440 Δ*phaABC* Δ*mvaB* Δ*hbdH* with plasmid pIY763 | This study |
| IY1101 | JBEI-233629 | *P. putida* KT2440 Δ*phaABC* Δ*mvaB* Δ*hbdH* Δ*ldhA* with plasmid pIY672 | This study |
| IY1102 | JBEI-233631 | *P. putida* KT2440 Δ*phaABC* Δ*mvaB* Δ*hbdH* Δ*ppsA* with plasmid pIY672 | This study |
| IY1200 | JBEI-233633 | *P. putida* KT2440 Δ*phaABC* Δ*mvaB* Δ*hbdH* Δ*PP_2675* with plasmid pIY672 | This study |
| IY1245 | JBEI-233635 | *P. putida* KT2440 WT with plasmid pIY670 | This study |
| IY1246 | JBEI-233637 | *P. putida* KT2440 Δ*PP_2675* with plasmid pIY670 | This study |
| IY1249 | JBEI-233639 | *P. putida* KT2440 Δ*phaABC* Δ*mvaB* with plasmid pIY670 | This study |
| IY1251 | JBEI-233641 | *P. putida* KT2440 Δ*phaABC* Δ*mvaB* Δ*gntZ* with plasmid pIY670 | This study |
| IY1252 | JBEI-233643 | *P. putida* KT2440 Δ*phaABC* Δ*mvaB* Δ*hbdH* Δ*aceA* with plasmid pIY670 | This study |
| IY1254 | JBEI-233645 | *P. putida* KT2440 Δ*phaABC* Δ*mvaB* Δ*hbdH* Δ*aceA* Δ*gntZ* with plasmid pIY670 | This study |
| IY1261 | JBEI-233647 | *P. putida* KT2440 Δ*phaABC* with plasmid pIY670 | This study |
| IY1262 | JBEI-233649 | *P. putida* KT2440 Δ*phaABC* Δ*mvaB* Δ*hbdH* Δ*ldhA* with plasmid pIY670 | This study |
| IY1263 | JBEI-233651 | *P. putida* KT2440 Δ*phaABC* Δ*mvaB* Δ*hbdH* Δ*ppsA* with plasmid pIY670 | This study |
| IY1319 | JBEI-233653 | *P. putida* KT2440 Δ*mvaB* with plasmid pIY670 | This study |

| | | | |
|---|---|---|---|
| IY1320 | JBEI-233655 | *P. putida* KT2440 Δ*hbdH* with plasmid pIY670 | This study |
| IY1452 | JBEI-233661 | *P. putida* KT2440 Δ*phaABC* Δ*mvaB* Δ*hbdH* Δ*ldhA* Δ*PP_2675* with plasmid pIY670 | This study |
| IY1884 | JBEI-233657 | *P. putida* KT2440 Δ*phaABC* Δ*PP_2675* with plasmid pIY670 | This study |
| IY1885 | JBEI-233659 | *P. putida* KT2440 Δ*phaABC* Δ*PP_2675* Δ*ldhA* with plasmid pIY670 | This study |

**Supplementary Table 2.** Plasmids used in this study

| Plasmids | Description | Reference |
|---|---|---|
| pXW1 | pBbB5k-$MvaS_{ef}$-$MvaE_{ef}$-T1-$MK_{mm}$-$PMD_{HKQ}$ | Wang et al. 2022 |
| pK18-*mvaB* | Plasmid to knockout *mvaB* (PP_3540) | This study |
| pK18-*aceA* | Plasmid to knockout *aceA* (PP_4116) | This study |
| pK18-*gntZ* | Plasmid to knockout *gntZ* (PP_4043) | This study |
| pK18-*hbdH* | Plasmid to knockout *hbdH* (PP_3073) | This study |
| pK18-*gltA* | Plasmid to knockout *gltA* (PP_4194) | This study |
| pK18-*ldhA* | Plasmid to knockout *ldhA* (PP_1649) | This study |
| pK18-*ppsA* | Plasmid to knockout *ppsA* (PP_2082) | This study |
| pIY554 | pRK2-Kan-*araC*-$P_{BAD}$-$MvaS_{ef}$-$MvaE_{ef}$-$T_{rpoH}$-$P_{trc1\text{-}O}$-$MK_{mm}$-$PMD_{HKQ}$ | This study |
| pIY602 | pBBR1-B5-Kan-*lacI*-$P_{lacUV5}$-$MvaS_{ef}$-$MvaE_{ef}$-$T_{rpoH}$-$P_{trc1\text{-}O}$-$MK_{mm}$-$PMD_{HKQ}$ | This study |
| pIY603 | pRK2-Kan-*lacI*-$P_{lacUV5}$-$MvaS_{ef}$-$MvaE_{ef}$-$T_{rpoH}$-$P_{trc1\text{-}O}$-$MK_{mm}$-$PMD_{HKQ}$ | This study |
| pIY604 | pRSF1010-Kan-*lacI*-$P_{lacUV5}$-$MvaS_{ef}$-$MvaE_{ef}$-$T_{rpoH}$-$P_{trc1\text{-}O}$-$MK_{mm}$-$PMD_{HKQ}$ | This study |
| pIY605 | pBBR1-Kan-*lacI*-$P_{lacUV5}$-$MvaS_{ef}$-$MvaE_{ef}$-$T_{rpoH}$-$P_{trc1\text{-}O}$-$MK_{mm}$-$PMD_{HKQ}$ | This study |
| pIY670 | pRK2-Kan-*araC*-$P_{BAD}$-$MvaS_{ef}$-$MvaE_{ef}$-$T_{rpoH}$-$P_{trc1\text{-}O}$-$MK_{mm}$-$PMD_{HKQ}$-AphA | This study |
| pIY671 | pRK2-Kan-*araC*-$P_{BAD}$-$MvaS_{ef}$-$MvaE_{ef}$-$T_{rpoH}$-$P_{trc1\text{-}O}$-$MK_{mm}$-$PMD_{HKQ}$-NudB | This study |
| pIY672 | pRK2-Kan-*araC*-$P_{BAD}$-$MvaS_{ef}$-$MvaE_{ef}$-$T_{rpoH}$-$P_{trc1\text{-}O}$-$MK_{mm}$-$PMD_{HKQ}$-AphA-NudB | This study |
| pIY697 | pRK2-Kan-*araC*-$P_{BAD}$-$MvaS_{ef}$-$MvaE_{ef}$-$T_{rpoH}$-$P_{trc1\text{-}O}$-$MK_{mm}$-$PMD_{HKQ}$-PhoA | This study |

| | | |
|---|---|---|
| pIY761 | pRK2-Kan-*araC*-$P_{BAD}$-$MvaS_{ef}$-$MvaE_{ef}$-AtoB-$T_{rpoH}$-$P_{trc1\text{-}O}$-$MK_{mm}$-$PMD_{HKQ}$-AphA-NudB | This study |
| pIY762 | pRK2-Kan-*araC*-$P_{BAD}$-$MvaS_{ef}$-$MvaE_{ef}$-NphT7-$T_{rpoH}$-$P_{trc1\text{-}O}$-$MK_{mm}$-$PMD_{HKQ}$-AphA-NudB | This study |
| pIY763 | pRK2-Kan-*araC*-$P_{BAD}$-$MvaS_{ef}$-$MvaE_{ef}$-PMK-$T_{rpoH}$-$P_{trc1\text{-}O}$-$MK_{mm}$-$PMD_{HKQ}$-AphA-NudB | This study |
| pIY765 | pRK2-Kan-*araC*-$P_{BAD}$-$MvaS_{ef}$-$MvaE_{ef}$-PMK-AtoB-$T_{rpoH}$-$P_{trc1\text{-}O}$-$MK_{mm}$-$PMD_{HKQ}$-AphA-NudB | This study |
| pIY853 | pK18-*PP_2675* | This study |

**Supplementary Table 3.** Minimal M9 medium recipe

10X M9 Salts

| Compound | Final concentrations |
|---|---|
| $Na_2HPO_4$ | 68 g |
| $KH_2PO_4$ | 30 g |
| NaCl | 5 g |

1X minimal M9 medium solution

| Compound/Stock | Per 1 L | Comments |
|---|---|---|
| 10X M9 Salts | 100 mL | Make 10x stock/filter separately |
| 1M $MgSO_4$ | 2 mL | Make 1 M solution/filter separately |
| 1M $CaCl_2$ | 100 µL | Make 1 M solution/filter separately |
| MQ $H_2O$ | 787.4 mL | Autoclaved |
| 20% Glucose | 100 ml | Autoclaved |
| Trace elements solution | 500 µL | Teknova |
| $(NH_4)_2SO_4$ | 10 ml | 1 M stock |
