## Supplementary material for "Genome-scale and pathway engineering for the sustainable aviation fuel precursor isoprenol production in *Pseudomonas putida*": SI_File_5_Combined_Scores

February 10, 2023

[1]:
```
import pandas as pd
import cobra
```

[2]:
```
model = cobra.io.load_json_model('iJN1463_IPP_bypass.json')
```

### 1 Load EMA-based and Opt-based targets

[3]:
```
EMA_score = pd.read_csv('EMA-based_targets.csv', index_col=0)
EMA_score.sort_values('EMA-based_Score', ascending=False).head()
```

[3]:

| | cMCS | EFM tool | EMA-based_Score |
|---|---|---|---|
| GLCDpp | 0.671667 | 0.0 | 0.636194 |
| PDH | 0.616111 | 0.0 | 0.583573 |
| LDH_D | 0.500000 | 1.0 | 0.526406 |
| ACACT1r | 0.497778 | 0.0 | 0.497778 |
| PPS | 0.450000 | 1.0 | 0.479047 |

[4]:
```
Opt_score = pd.read_csv('Opt-based_targets.csv', index_col=0)
Opt_score.sort_values('Opt-based_Score', ascending=False).head()
```

[4]:

| | OptKnock | OptKnock_blocked_exchanges | Opt-based_Score |
|---|---|---|---|
| ACt2rpp | 0.958333 | 0.885932 | 0.922132 |
| CS | 0.758333 | 0.908745 | 0.833539 |
| HMGL | 0.958333 | 0.437262 | 0.697798 |
| BDH | 0.966667 | 0.395437 | 0.681052 |
| TPI | 0.950000 | 0.403042 | 0.676521 |

[5]:
```
Combined = EMA_score[['EMA-based_Score']].join(Opt_score['Opt-based_Score'],␣
↪how='outer')
Combined.head()
```

[5]:

| | EMA-based_Score | Opt-based_Score |
|---|---|---|
| 1PPDCRc | 0.011111 | 0.001901 |
| 3HAACOAT100 | 0.055556 | NaN |
| 3HAACOAT120 | 0.055556 | NaN |
| 3HAACOAT121 | 0.055556 | 0.004167 |
| 3HAACOAT140 | NaN | 0.007969 |

### 2 Filter essential reactions, transport reactions, and reactions without genes

[6]:
```
gene_essentiality = pd.read_table('Gene_Essentiality.txt', index_col=0)
gene_essentiality.head()
```

[6]:
```
           Proteins   RbTnSeq Data
Genes
PP_s0001  PP_s0001    spontaneous
PP_3591       dpkA  Non Essential
PP_1257    PP_1257  Non Essential
PP_2136       fadB  Non Essential
PP_2008       fadH  Non Essential
```

[7]:
```
essential_reactions = [x for x in Combined.index if ((x in model.reactions) and
                        any(gene_essentiality.loc[g.id, 'RbTnSeq Data'] ==␣
 ↪'Essential' for g in
                            model.reactions.get_by_id(x).genes if g.id in␣
 ↪gene_essentiality.index))]
Combined.drop(essential_reactions, inplace=True)
```

[8]:
```
transporters = [x for x in Combined.index if ((x in model.reactions) and
                (all(c in model.reactions.get_by_id(x).compartments for c in␣
 ↪['c','p']) or
                 all(c in model.reactions.get_by_id(x).compartments for c in␣
 ↪['e','p'])))]
Combined.drop(transporters, inplace=True)
```

[9]:
```
reactions_without_genes = [x for x in Combined.index if ((x in model.reactions)␣
 ↪and
                            not model.reactions.get_by_id(x).genes)]
Combined.drop(reactions_without_genes, inplace=True)
```

### 3 Rank targets for each method and choose top priority targets

[10]:
```
Combined_rank = Combined.rank(ascending=False)
Combined_rank['Combined_rank'] = Combined_rank.mean(axis=1, skipna=False).rank()
Combined_rank.sort_values(by='Combined_rank', inplace=True)
```

**Top 10 targets by both EMA-based and Opt-based methods**

[11]:
```
Combined_rank_top10 = Combined_rank[(Combined_rank['EMA-based_Score'] <= 10) &␣
 ↪(Combined_rank['Opt-based_Score'] <= 10)]
Combined_rank_top10
```

[11]:

| | EMA-based_Score | Opt-based_Score | Combined_rank |
|---|---|---|---|
| HMGL | 9.0 | 2.0 | 1.5 |
| GND | 5.0 | 6.0 | 1.5 |
| BDH | 9.0 | 3.0 | 3.0 |

**Top targets by EMA-based method, but not top 10 by Opt-based method**

[12]:
```
Combined_rank.sort_values(by='EMA-based_Score').drop(Combined_rank_top10.
↪index)[0:5]
```

[12]:

| | EMA-based_Score | Opt-based_Score | Combined_rank |
|---|---|---|---|
| LDH_D | 1.0 | NaN | NaN |
| ACACT1r | 2.0 | 21.0 | 6.0 |
| PPS | 3.0 | 91.0 | 28.0 |
| KAT1 | 4.0 | 26.0 | 10.0 |
| PC | 6.0 | 44.0 | 16.0 |

**Top targets by Opt-based method, but not top 10 by EMA-based method**

[13]:
```
Combined_rank.sort_values(by='Opt-based_Score').drop(Combined_rank_top10.
↪index)[0:5]
```

[13]:

| | EMA-based_Score | Opt-based_Score | Combined_rank |
|---|---|---|---|
| CS | 19.0 | 1.0 | 5.0 |
| TPI | 21.0 | 4.0 | 7.5 |
| PGK | 12.0 | 5.0 | 4.0 |
| GARFT | 34.0 | 7.0 | 15.0 |
| ICL | 57.0 | 8.0 | 18.0 |
